# Molecular Recognition of Itching Neuropeptides by Bombesin Receptors

**DOI:** 10.1101/2022.10.10.511544

**Authors:** Changyao Li, Youwei Xu, Heng Liu, Hongmin Cai, Yi Jiang, H. Eric Xu, Wanchao Yin

**Affiliations:** The CAS Key Laboratory of Receptor Research, Shanghai Institute of Materia Medica, Chinese Academy of Sciences, Shanghai 201203, China; University of Chinese Academy of Sciences, Beijing 100049, China; Lingang Laboratory, Shanghai 200031, China; School of Life Science and Technology, ShanghaiTech University, Shanghai 201210, China; Zhongshan Institute for Drug Discovery, Shanghai Institute of Materia Medica, Chinese Academy of Sciences, Guangdong 528400, China

**Author notes:** Correspondence (H.E.X), (W.Y.). These authors contributed equally (Changyao Li, Youwei Xu, Heng Liu).

## Abstract

Neuromedin B (NMB) and gastrin-releasing peptide (GRP), two bombesin analogs, are endogenous itch-specific neuropeptides that induce histaminergic and nonhistaminergic itch, respectively. Their functions are mediated by two G protein-coupled bombesin receptors, NMBR and GRPR. Here we present cryo-electron microscopy(cryo-EM) structures of G protein coupled NMBR and GRPR bound to NMB and GRP, respectively. The structures reveal that both bombesin receptors contain an extended and deep pocket to adopt NMB and GRP, with the conserved C-terminal motif of GH(F/L)M from both peptides to contact the toggle switch residues for the receptor activation. Together with mutational and functional data, our structures reveal the mechanism of ligand selectivity and specific activation of the bombesin receptors. These findings also pave the way to facilitate rational design of therapies targeting bombesin receptors for the treatment of pruritus.

## INTRODUCTION

Itch, as one of the most primal sensations, is an essentially protective mechanism to remind animals of impairment and expel noxious environmental substances, but often exists in patients with allergic skin diseases and/or systemic and neuropathic problems. On occasions, persistent itch (or chronic pruritus) significantly affects sleep, mood and individual health, and reduces life quality of patients. Unfortunately, there is no FDA-approved treatment for chronic pruritus despite great efforts on antipruritic research in the past few decades. Depending on whether histamine release is involved, itch can be divided into two subtypes: histaminergic itch (evoked by histamine, serotonin (5-HT), compound 48/80, and defensin) and nonhistaminergic itch (evoked by chloroquine, SLIGRL, and BAM8-22)^1^. Several potential receptors for pruritus perception have been investigated, including the histaminergic receptors H_1_R^2^, and the recently reported nonhistaminergic receptors MRGPRX2 and MRGPRX4^3,4^, setting the stage for future efforts to endogenous ligands for strategies to fight itch. Although almost all itch arises from primary sensory neurons innervating the skin, itch signals must ultimately enter the spinal cord through dorsal root ganglia (DRG) neurons to activate a circuit of spinal interneurons, which finally evoke a perception of itch or scratching reflex. Many molecular targets, including ion channels and G-protein-coupled receptors (GPCRs), are involved in mediating and sensing pruritus^5^. Therefore, together with the primary sensory pruriceptors as mentioned above, the itch-specific neuropeptide receptors in the spinal cord are at the core of itch transmission. Gastrin-releasing peptide(GRP) and gastrin-releasing peptide receptor (GRPR or BB_2_ receptor) are among the first neuropeptide-receptor pairs identified to constitute the itch-specific pathway in the nervous system^6^. The roles of GRP and GRPR in itch transmission have shed light on itch perception and signal transduction, and established GRPR as a tractable target for therapeutic intervention.

Bombesin (Bn) is an amphibian tetradecapeptide firstly found in the skin of *European Bombina*. Two mammalian Bn-like peptides have been characterized, neuromedin B (NMB) and GRP, which are the endogenous ligands for neuromedin B receptor (NMBR or BB_1_ receptor) and GRPR, respectively^7^. In addition, there is an orphan bombesin receptor, BRS3 or BB_3_R, which shares sequence and structural similarity with NMBR and GRPR but is not activated by NMR or GRP (Supplementary information, Fig. S1)^7^. Both NMR and GRP are widely distributed in mammals especially in the central nervous system (CNS) and gastrointestinal tract, and they are involved in a variety of physiological and pathological processes, including contraction of smooth muscle, exocrine and endocrine secretion, and multiple cancers. Most importantly, increasing studies have revealed that NMBR and GRPR integrate itch-specific pathways in the nervous system, mediating histaminergic and nonhistaminergic itches^1,5,6,8-10^, respectively (Fig. 1a). As itch-specific neuropeptide receptors in the spinal cord, both NMBR and GRPR play a pivotal role in itch biology; accordingly, they are attractive targets for antipruritic intervention. Extensive efforts have been made to elucidate the mechanisms of agonism or antagonism at bombesin receptors through mutagenesis studies^11-14^, but the lack of precise structural information largely impedes our understanding of the molecular details of ligand recognition and receptor activation. In this study, we determined the structures of NMBR and GRPR, both complexed with the engineered G_q_ and selective peptide agonists NMB30 and GRP (14-27) at global resolutions of 3.15 Å and 3.30 Å, respectively, using single-particle cryo-EM. Combined with functional data, the structures provide the molecular basis of peptide recognition by the itch receptors and set for understanding itch perception. The structures also provide a basis for rational drug design for pruritus treatment by targeting NMBR and GRPR.

**Fig. 1.**
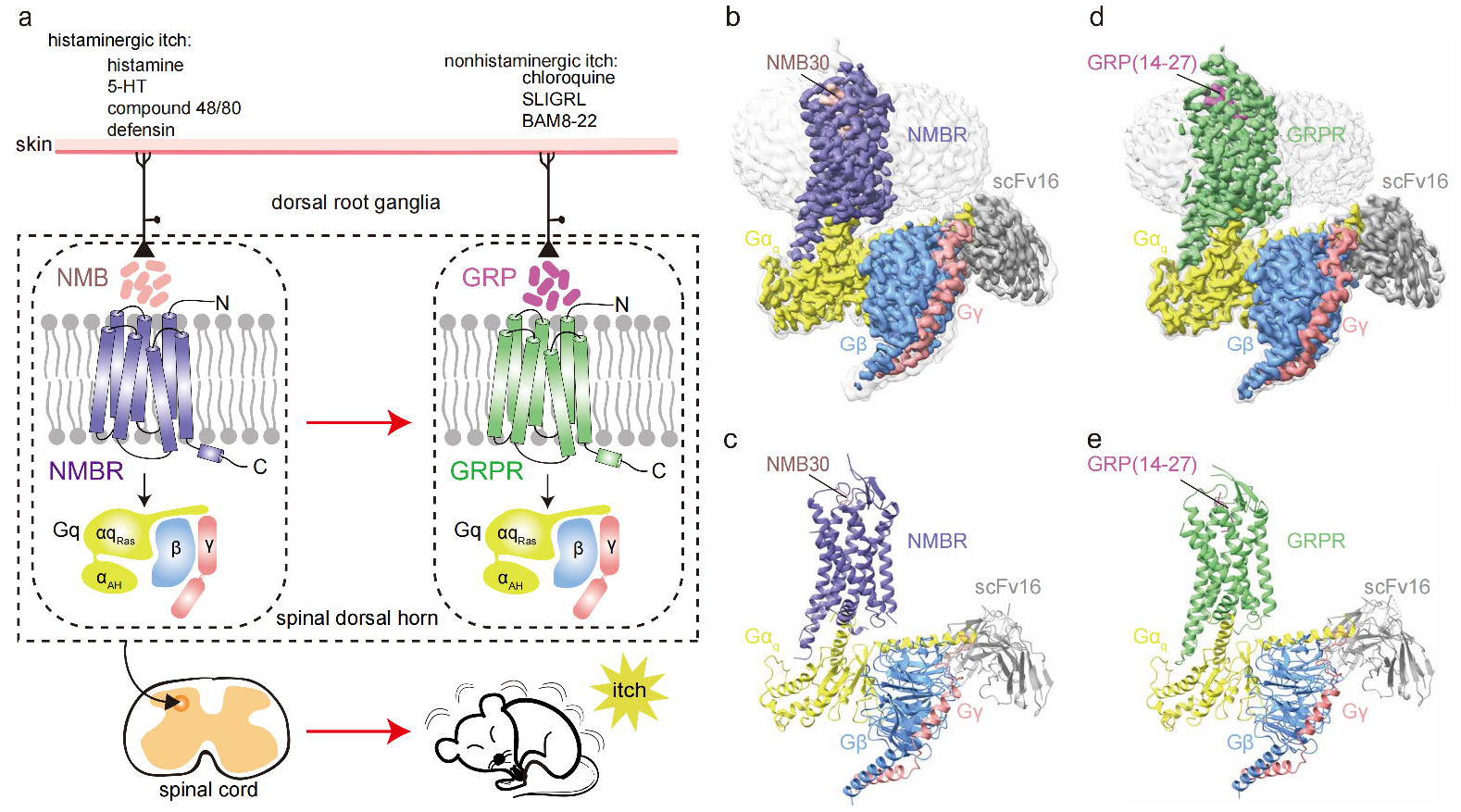
Cryo-EM structures of the NMB30-NMBR-G_q_ and GRP (14-27)-GRPR-G_q_ complexes. **a** Schematic illustration of the neuropeptide coding in the chemical itch pathways. Briefly, itch induced by histaminergic or nonhistaminergic pruritogens is encoded by either NMB or GRP, which would activate their receptors NMBR and GRPR in the dorsal root ganglia (DRG) neurons, respectively. The signal transmission in both GRPR and NMBR neurons is mediated through the heterotrimeric G_q_ proteins. GRPR neurons could integrate nonhistaminergic itch, together with the histaminergic itch information from NMBR neurons. Finally, the itch signal is further transmitted to dorsal horn of spinal cord, finally to brain that ultimately causes itch sensation. **b-c** Orthogonal views of the density map (**b**) and model (**c**) for the NMB30-NMBR-G_q_ complex. NMB30 is shown in coral; the receptor NMBR is displayed in slate blue. The heterotrimeric G_q_ proteins are colored by subunits: Gα_q_, yellow; Gβ, cornflower blue; Gγ, deep salmon; scFv16, gray. **d-e** Orthogonal views of the density map (**d**) and model (**e**) of the GRP (14-27)-GRPR-G_q_ complex. GRP (14-27) is shown in orchid; the receptor GRPR is displayed in green. The heterotrimeric G_q_ proteins are colored by subunits: Gα_q_, yellow; Gβ, cornflower blue; Gγ, deep salmon; scFv16, gray.

## RESULTS

### Structure determination of NMB30-NMBR-G_q_ and GRP (14-27)-GRPR-G_q_ complexes

We fused thermo-stabilized fusion protein BRIL at the N terminus of both NMBR and GRPR, and applied the NanoBiT tethering strategy to improve complex stability and homogeneity^15,16^ (Supplementary information, Fig. S2a, d). To facilitate the cryo-EM studies, we use an engineered G_q_ protein chimera, which was generated on the basis of the miniGα_s/q_71 scaffold with the N terminus (residues 1-18) replaced by the N terminus (residues 1-18) of G_i_ protein to render the binding ability to the scFv16 antibody^17,18^. Analogous approaches with BRIL fusion, NanoBiT tethering, and G_q_ engineering had previously been used successfully in solving several GPCR/G-protein complexes^19,20^. Unless otherwise specified, G_q_ refers to engineered G_q_ protein chimera, which is used for further structure studies. We also introduced the point mutation of Q^3.43^ into NMBR and GRPR, inspired by the common activation mechanism of class A GPCRs^21^ (L134^3.43^ for NMBR and L131^3.43^ for GRPR, superscripts refer to Ballesteros–Weinstein numbering^22^). We prepared both complexes by co-expression of the receptors with Gα_q_ and Gβγ, in the presence of corresponding endogenous agonist (NMB30 for NMBR and GRP (14-27) for GRPR), and the antibody scFv16, which allowed the efficient assembly and further stabilization of agonist-bombesin receptor-G_q_-scFv16 complexes (Supplementary information, Fig. S2b, c, e and f).

The structure of the NMB30-NMBR-G_q_ complex was defined with 355,509 final particles from 3,034,736 initial particles to a global nominal resolution of 3.15 Å (Supplementary information, Fig. S3 and Table S1). The structure of the GRP (14-27)-GRPR-G_q_ complex was defined with 88,357 final particles from 3,365,839 initial particles to a global nominal resolution of 3.30 Å (Supplementary information, Fig. S4 and Table S1). The cryo-EM maps are sufficiently clear to define the position of the receptor, the G-protein trimer, the scFv16, and bound ligand in both receptor–G-protein complexes, with the majority of amino acid side chains resolved in the final refined maps (Fig. 1b-e; Supplementary information, Figs. S3-S5). Both complexes display highly similar overall structures with the root mean square deviation (RMSD) of 0.7 Å for the Cα atoms of the entire complexes (Supplementary information, Fig. S6a) and 0.8 Å for the Cα atoms between NMBR and GRPR (Supplementary information, Fig. S6b).

### Recognition of NMB30 by NMBR

Unambiguous electron densities were observed for the C-terminal 9 residues (NLWATGHFM-NH_2_) in NMB30, while the long N-terminal part of NMB30 was not modeled because of the poor density map. The visibility of the C-terminal 9-residue peptide is close to the minimal 10-residue peptide required for full potency equal to NMB30^23^. NMB30 occupies the orthosteric binding pocket comprising all three extracellular loops (ECLs) and most TM helices, including TM2 to TM7 (Fig. 1b, c). It is interesting that NMB30 has a dumbbell-shaped overall conformation and adopts a binding pose perpendicular to the membrane plane (Fig. 2a), with its C terminus inserted deep into the transmembrane domain (TMD) bundle, while its N terminus (referring throughout the paper to the resolved N-terminal amino acids) points to the extracellular surface (Fig. 2a, b). The region of ligand recognition of NMB30 can be divided into two major parts: (1) the extracellular loops around the N-terminal dumbbell end; and (2) the bottom of the TMD pocket burying the C-terminal dumbbell end (Fig. 2b-e).

**Fig. 2.**
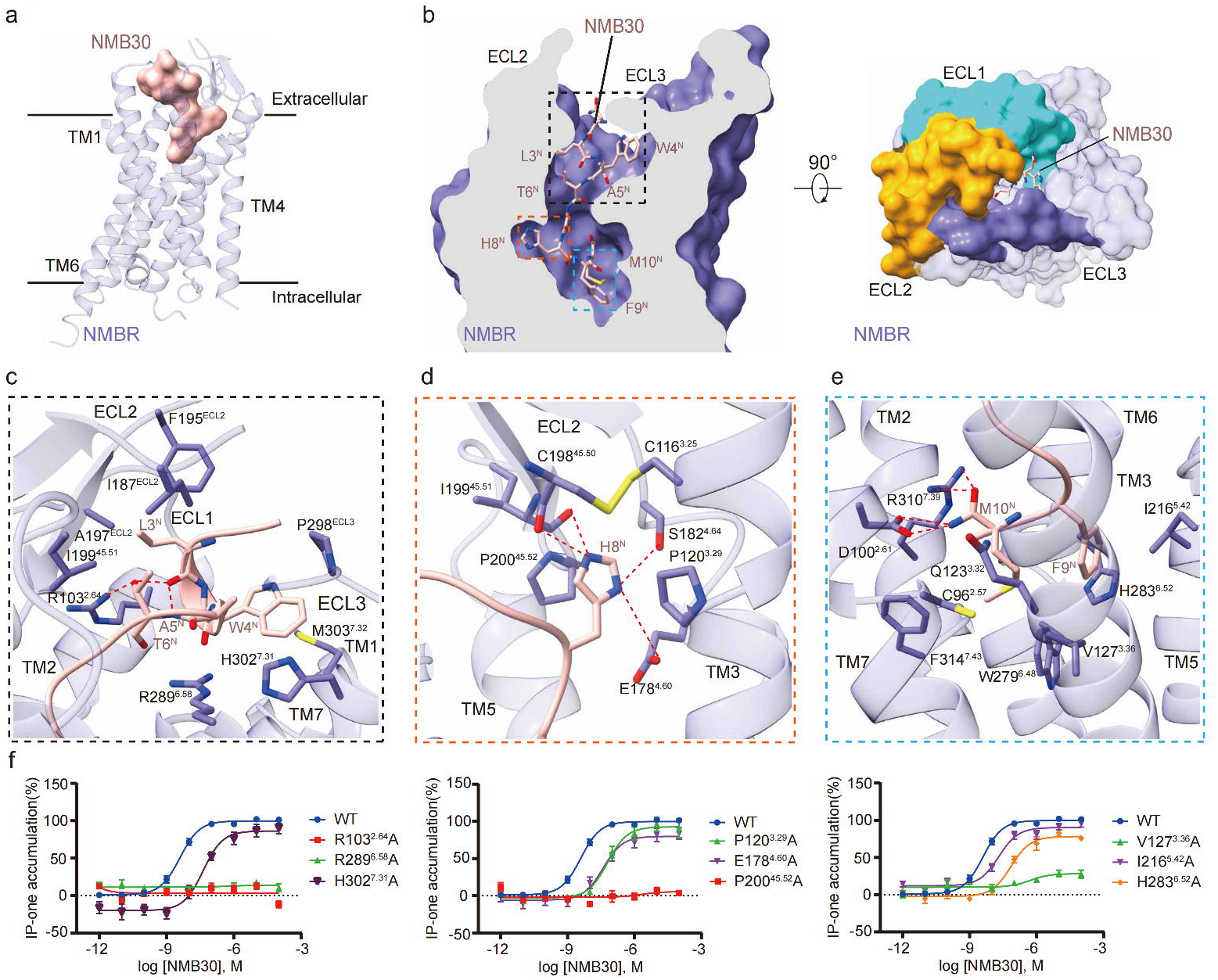
The NMB30-binding pocket in NMBR. **a** Side view of the NMB30 bound NMBR complex. The receptor is shown in slate blue cartoon representation. The agonist NMB30, shown with coral surface, presents a dumbbell-shaped overall conformation and adopts a binding pose perpendicular to the membrane plane. **b** Cross-section and top views of the NMB30-binding pocket in NMBR. Side chains of NMB30 residues are displayed as sticks. The ECLs from NMBR are annotated. **c-e** Detailed interactions of NMB30 with NMBR. The binding sites of L3^N^-T6^N^ (**c**), H8^N^ (**d**) and F9^N^ and M10^N^ (**e**) are shown. Hydrogen bonds are depicted as red dashed lines. **f** Effects of mutations in the NMB30-binding pocket in IP1 assay. Data are presented as mean ±S.E.M. of at least three independent experiments. Source data are available online.

At the extracellular side, three ECLs are folded to embrace the resolved N-terminal amino acids of NMB30, which looks like a dumbbell end. In order to illustrate the conservation of NMB30, GRP (14-27), and related naturally occurring peptides of the bombesin family, all positions of bombesin peptides are numbered from the amino terminus of NMB or NMC for clarity (Supplementary information, Table S2). The N-terminal dumbbell end comprises NLWAT (2-6)^N^ (The superscript N refers to NMB30) sequences, which fold into a short α-helix (Fig. 2c). At one side of the helix, the side chains of L3^N^ and T6^N^ are buried in a hydrophobic cavity formed by the residues of I187^ECL2^, F195^ECL2^, A197^ECL2^, and I199^45.51^ (Fig. 2c). In addition, the side chain of T6^N^ forms a close hydrogen bond with R103^2.64^ from TM2 (Fig. 2c), which further stabilizes the conformation of the N-terminal dumbbell end and plays a vital role in NMBR activation. This is consistent with our mutation assay that the R103^2.64^A mutation completely inactivates the NMBR (Fig. 2f; Supplementary information, Fig. S7a-e and Table S3). At the opposite side of the helix, the side chain of W4^N^ is held by hydrophobic interactions with residues P298^ECL3^, H302^7.31^, and M303^7.32^ (Fig. 2c), of which alanine substitution of H302^7.31^ decreases the potency of NMB30 to activate NMBR (Fig. 2f; Supplementary information, Fig. S7a-e and Table S3). Presumably, all these interactions restrict the orientation of the N terminus of NMB30 toward the extracellular solvent.

At the bottom of the binding pocket is the C-terminal dumbbell end formed by the HFM motif of NMB30 (Fig. 2b, d and e). The side chain of H8^N^ is well sandwiched by the P120^3.29^ from TM3 and P200^45.52^ from ECL2, which is further stabilized by the conserved disulfide bond between C116^3.25^ and C198^45.50^ (Fig. 2d). In addition, the side chain of H8^N^ also forms two hydrogen bonds with side chains of E178^4.60^ and S182^4.64^, and two additional hydrogen bonds with backbone carbonyl groups of C198^45.50^ and I199^45.51^ (Fig. 2d). This binding mode is supported by our mutagenesis assays, in which alanine mutations of P120^3.29^, E178^4.60^, and P200^45.52^ had a great influence on the receptor activation, especially for the P200^45.52^A and P200^45.52^S mutations, suggesting the key role of ECL2 in peptide recognition (Fig. 2f; Supplementary information, Fig. S7a-e and Table S3). Additionally, the H8^N^ plays a pivotal role in binding to NMBR, as reported that the synthetic bombesin-related peptide Roh-Lit bound to GRPR with approximately 1000-fold higher affinity than LeuPLL, of which the equivalent H8 was replaced by S8 (Supplementary information, Table S2)^24^. The penultimate F9^N^ forms a stabilizing π-π packing with H283^6.52^, which is supported by mutagenesis studies that alanine replacement of H283^6.52^ associated with about 20-fold reduction in NMB30 potency in inducing IP production (Fig. 2f; Supplementary information, Fig. S7a-e and Table S3). In addition, the side chains of F9^N^ and M10^N^ fit into a hydrophobic crevice formed by the residues of C96^2.57^, V127^3.36^, W279^6.48^, H283^6.52^, and F314^7.43^ (Fig. 2e). The hydrophobic network here is essential for NMBR activation, since substituting V127^3.36^ with alanine entirely abolished the activity of NMB30 (Fig. 2f; Supplementary information, Fig. S7a-e and Table S3). It should be noted that, with the assistance of hydrophobic network, the C-terminal amidated end turns a large angle back to nearly parallel with the main chain of ATGH sequence of NMB30, and forms extensive polar interactions with the residues from TM2, TM3 and TM7 (Fig. 2e). The C-terminal amide makes a hydrogen bond with the side chain of D100^2.61^, while the C-terminal carbonyl group forms a hydrogen bond with the side chain of neighboring R310^7.39^ (Fig. 2e), which are both critical to NMBR activation. Alanine replacements of D100^2.61^ and R310^7.39^ displayed a much more predominant influence showing a 10- to 50-fold reduction in NMB30 potency in inducing receptor activation (Supplementary information, Fig. S7a-e and Table S3). It is noteworthy that the remarkable conformational shift of the C-terminal amidated end plays a pivotal role in NMB30 potency in inducing receptor activation, since our mutation of Q123^3.32^ to alanine only associated with about 30-fold reduction in NMB30 potency, while the mutation of Q123^3.32^ to arginine with a bulky side chain to clash with the C-terminal amidated end completely abolished the ability of NMB30 to stimulate NMBR (Fig. 2e, f; Supplementary information, Fig. S7a-e and Table S3). Together, these detailed structural analyses provide important information to better understand the recognition mechanism of NMB30 by NMBR.

### Recognition of GRP (14-27) by GRPR

The structure of the GRP (14-27)-GRPR-G_q_ complex also revealed the clear density for the C-terminal 9 residues (NHWAVGHLM) of GRP (Fig. 1d, e). Similar to the NMB30 structure, the GRP (14-27) structure also exhibits a dumbbell-shaped conformation (Fig. 3a). Inspection into the detailed interactions between GRP (14-27) and GRPR shows a similar orthosteric binding pocket in GRPR (Fig. 1d, e), partially explaining the conserved activation mechanism between these two bombesin receptors. In order to facilitate elaboration, we also divided the region of ligand recognition of GRP (14-27) into two major parts, as in the NMB30 structure: (1) the extracellular loops around the N-terminal dumbbell end; and (2) the bottom of the TMD pocket burying the C-terminal dumbbell end. Nevertheless, distinct interactions are observed between two peptides and the corresponding receptor subtypes, providing the basis for their receptor selectivity as described below (Fig. 3b-e).

**Fig. 3.**
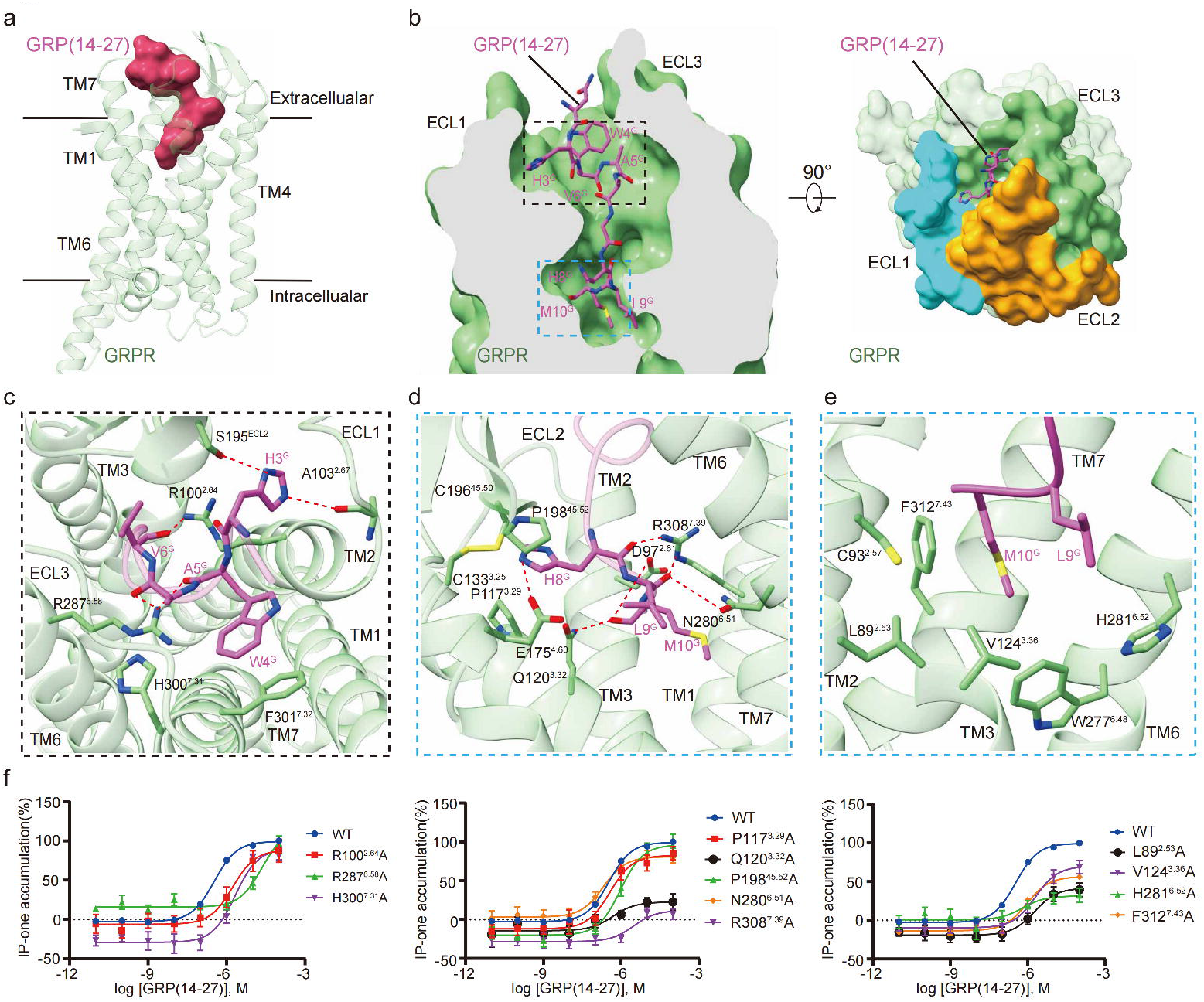
The GRP (14-27)-binding pocket in GRPR. **a** Side view of the GRP (14-27) bound GRPR complex. The receptor is shown in green cartoon representation. GRP (14-27) with a dumbbell-shaped overall conformation is shown with orchid surface. **b** Cross-section and top views of the GRP (14-27)-binding pocket in GRPR. Side chains of the GRP (14-27) residues are displayed as sticks. The ECLs from GRPR are colored differently: ECL1, in cyan; ECL2 in orange and ECL3 in blue. Detailed interactions of GRP (14-27) with GRPR. The binding sites of H3^G^-V6^G^ (**c**) and H8^G^-M10^G^ (**d-e**) are shown, with hydrogen bonds depicted in red dashed lines. **f** Effects of mutations in the GRP (14-27)-binding pocket in IP1 assay. Data are presented as mean ±S.E.M. of at least three independent experiments. Source data are available online.

The resolved NHWAV (2-6)^G^ (The superscript G refer to GRP (14-27)) sequences in GRPR, resembling the N-terminal dumbbell end, were also embraced by the squashed cavity folded by the three ECLs (Fig. 3b, c). Compared with the NMB30 structure, there is no strong intramolecular hydrogen bond in GRP (14-27) and the resolved N-terminal amino acids adopt a more extended conformation. The side chains of R100^2.64^ and R287^6.58^ make three hydrogen bonds with the main-chain carbonyl groups of W4^G^, A5^G^, and V6^G^, and drag the N terminus of GRP (14-27) toward ECL1 and ECL2, where the side chain of H3^G^ forms a hydrogen bond with the backbone carbonyl of A103^2.67^ (Fig. 3c). In addition, the side chain of W4^G^ forms aromatic packing with H300^7.31^ and F301^7.32^ (Fig. 3c), and further stabilizes the N-terminal bending conformation.

At the bottom of the binding pocket is the C-terminal dumbbell end formed by the HLM motif of GRP (14-27). Similar to the NMB30 structure (Fig. 3a, b), the 3 residues of the HLM sequence shape the C-terminal dumbbell end. H8^G^ is also sandwiched by the P117^3.29^ from TM3 and P198^45.52^ from ECL2, which is further stabilized by the conserved disulfide bond between C113^3.25^ and C196^45.50^, and the hydrogen bond between H8^G^ and the side chain of E175^4.60^ from TM4 (Fig. 3d). Additionally, R308^7.39^ forms two hydrogen bonds with the backbone carbonyl groups of H8^G^ and L9^G^, N280^6.51^ forms a hydrogen bond with the backbone carbonyl of L9^G^, and the C-terminal carbonyl of M10^G^ forms two hydrogen bonds with the side chains of D97^2.61^ and Q120^3.32^ (Fig. 3d). Besides, the side chains of L9^G^ and M10^G^ fit into a hydrophobic crevice formed by the residues of L89^2.53^, C93^2.57^, V124^3.36^, W277^6.48^, H281^6.52^, and F312^7.43^ (Fig. 3d). The hydrophobic network here is essential for GRPR activation, since alanine mutations of V124^3.36^, H281^6.52^, and F312^7.43^ displayed a predominant influence showing a 3- to 6-fold reduction in GRP (14-27) potency in inducing receptor activation (Fig. 3e; Supplementary information, Fig. S7f-j and Table S3). Together, these results reveal the recognition mechanism of GRP (14-27) and highlight the distinct peptide-binding modes between the two bombesin receptors.

### Basis for ligand selectivity of Bombesin Receptors

NMBR and GRPR have high affinity and selectivity for their respective endogenous agonists NMB30 and GRP (14-27). NMBR binds to NMB30 with 640-fold higher affinity than GRP(14-27), whereas GRPR binds to GRP(14-27) with 650-fold higher affinity than NMB30^7^. In terms of their sequence and structural features, NMB30 belongs to the ranatensin group^7^, which have a carboxyl terminus of GHFM-NH_2_ (Supplementary information, Table S2); GRP (14-27) belongs to the bombesin group^7^, which all have a carboxyl terminus of GHLM-NH_2_ (Supplementary information, Table S2). For both NMB30 and GRP (14-27), peptide length required for full potency is the C-terminal decapeptide^23^, which is named as NMB for NMB30, and named as NMC for GRP (14-27) (Supplementary information, Table S2). NMB30 differs from GRP (14-27) at three residues within the C-terminal decapeptide, and the mutations of L3^N^ and F9^N^ in NMB30 to the equivalent H3^G^ and L9^G^ in GRP (14-27) are known to increase potency and specificity^23^. Structural insights into the binding pockets encompassing the two peptides reveal that minor amino acid differences between the two receptors reshape their binding modes and allow different peptide preferences, whereas most residues in the binding pockets of NMBR and GRPR are highly conserved.

As previously described, the resolved lengths of NMB30 and GRP (14-27) converge in an approximately overlapped position and the residues interacting with the C-terminal residues of the two respective peptides in the receptors are similar (Fig. 4). The main differences are located in the ECL2 with a deflection of 2.1 Å as measured between the Cα atoms of the conserved F195^ECL2^ in NMBR and F193 ^ECL2^ in GRPR (Fig. 5a). Both side chains of residues at position 3 of the two bombesin peptides are docked into the cavity between ECL1 and ECL2(Fig. 5b). In addition, the swing of the side chain of F193^ECL2^ in GRPR away from the peptide GRP (14-27) creates a sufficient space to fit the bulky side chain of H3^G^, which is reported to be the most important residue for receptor subtype selectivity^23^. The characteristic conformation of F193 ^ECL2^ in GRPR is stabilized by the side chain of F184^ECL2^, which is shifted toward the A197^45.51^. The corresponding NMBR residues are I187^ECL2^ and I199^45.51^, which side chains are too rigid to allow similar ECL2 movement as seen in the GRPR structure. The NMBR ECL2 is further stabilized by a unique hydrogen bond from the backbone amide group of T196^ECL2^ to the side chain of E108^23.49^ (the cognate residue R105^23.49^ in GRPR pointing to the solvent). The locked conformation of F195^ECL2^ in NMBR greatly affects the affinity and activity of bombesin peptides with H3^G^ residues at position 3 of GRP (14-27), consistent with a detailed study of loss- and gain-of-affinity chimeric GRPR/NMBR^14^. In contrast, the ECL2 flexibility in GRPR allows its compatibility for L3 or Q3 in its peptide ligands, consistent with systematical studies of 24 peptide ligands at GRPR^24^. Therefore, our structures partially explain why NMBR binds to GRP (14-27) with very weak affinity in contrast to NMB30.

**Fig. 4.**
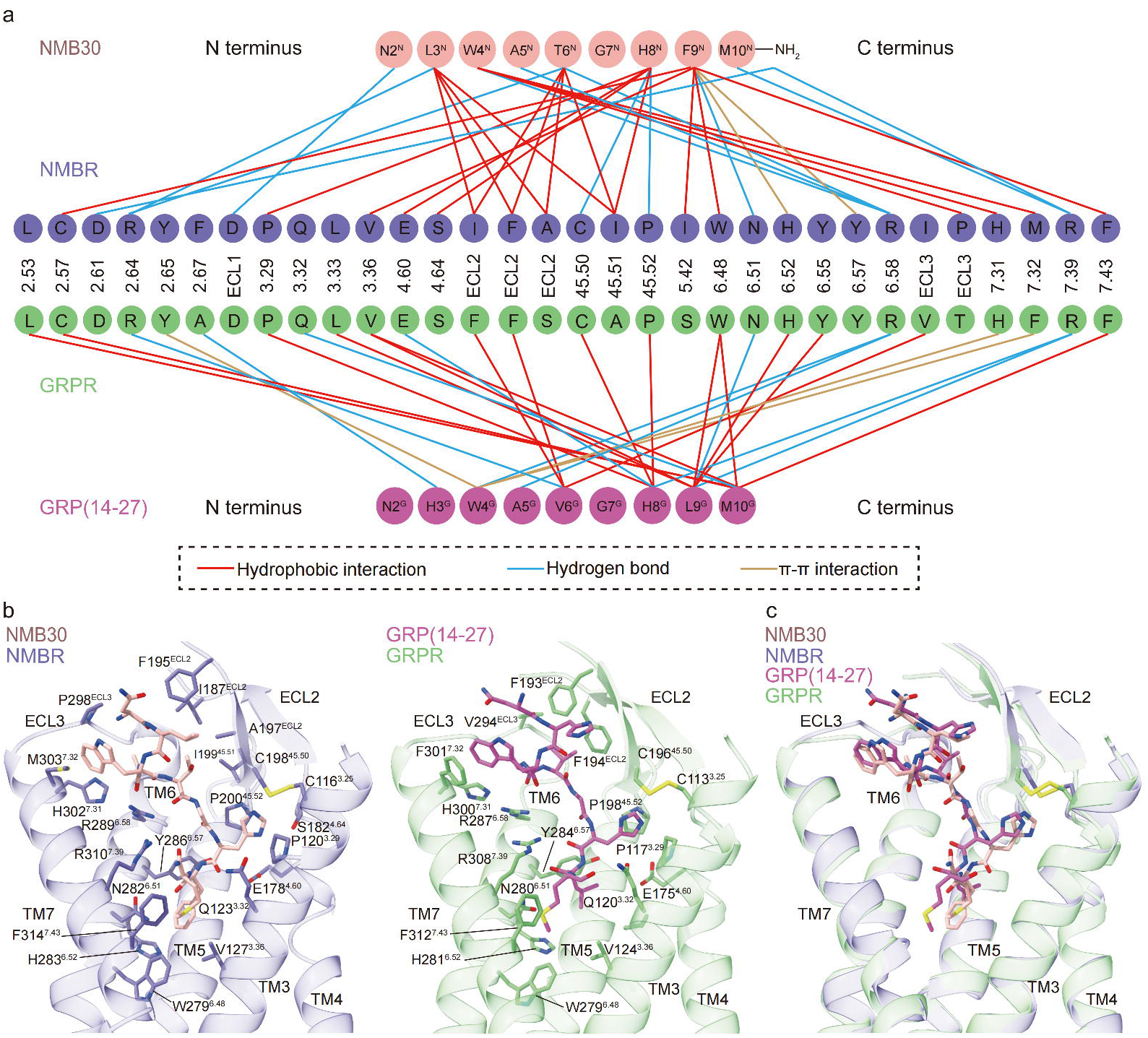
Comparison of interactions between NMB30 and GRP (14-27) with two bombesin receptors. **a** Representative interaction network of NMB30 bound to NMBR and GRP (14-27) bound to GRPR. Amino acids in NMB30 and GRP (14-27), as well as residues in corresponding binding pockets of NMBR and GRPR, are displayed as circled one-letter codes. Lines show interactions between peptides and bombesin receptor subtypes. **b** The binding pockets of NMB30 and GRP (14-27) in receptors NMBR and GRPR. **c** Superposition of the binding pockets of NMB30 and GRP (14-27). For (**b**) and (**c**), the ligands and receptors are presented as sticks and cartoon, respectively.

**Fig. 5.**
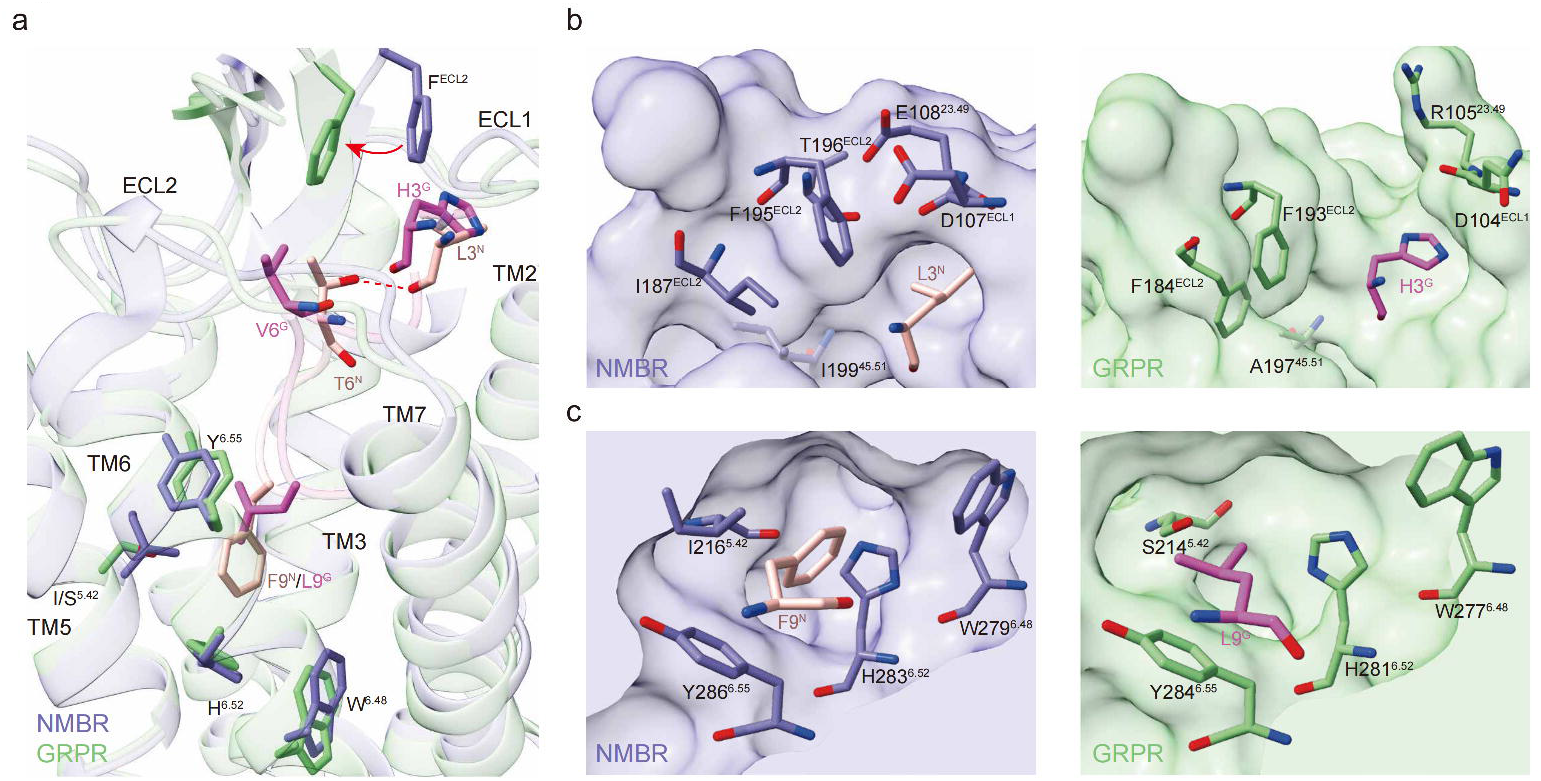
Molecular basis of bombesin peptides selectivity for bombesin receptors. **a** Detailed interactions between L3^N^ and F9^N^ with residues in NMBR, and H3^G^ and L9^G^ with residues in GRPR. **b** A comparison of detailed interactions of L3^N^ andH3^G^ and corresponding residues in NMBR and GRPR. **c** A comparison of detailed interactions of F9^N^ and L9^G^ and corresponding residues in NMBR and GRPR. For **a-c**, the related residues in the ligands and receptors are shown as sticks.

The selectivity of NMBR for NMB30 is also contributed by specific interactions of F9^N^ of NMB30 with conserved residues of W^6.48^, H^6.52^, and Y^6.55^, which forms a relatively hydrophobic binding pocket in both receptors (Fig. 5c). This NMBR pocket also contains I216^5.42^, which is directly packed against F9^N^ (Fig. 5c). The corresponding GRPR residue is S214^5.42^ (Fig. 5c), which cannot form the similar packing interactions with F9^N^ of NMB30, providing further explanation for NMBR selectivity for NMB30, which is consistent with previous studies with NMBR/GRPR chimeric receptors^11^. In addition, T6^N^ in NMB30 forms an intramolecular hydrogen bond interaction with L3^N^, which is absent in the structure of GRP (14-27) and has not been previously reported (Fig. 5a). The [V^6^] NMB showed approximately a 11-fold decrease in binding affinity to NMBR^23^, suggesting the hypothesis that the conformation of NMB30 assisted by this unique intramolecular hydrogen bond interaction also plays a role in receptor subtype selectivity. Together, these data reveal the determinants of bombesin receptor selectivity between NMB30 and GRP (14-27).

### G_q_ coupling mechanism of Bombesin Receptors

The overall structures of the NMBR and GRPR signaling complexes are similar to each other, with the overall engagement of G_q_ to both bombesin receptors mainly maintained by key interactions from ICL2, ICL3, TM5, TM6, and TM7-H8 kink regions (Fig. 6a, c). Compared to G_s_ complex, such as active β_2_AR (PDB 3SN6) both structures featured a less-pronounced outward swing movement of the intracellular tip of TM6, which is common to solved structures of other G_q_-coupled GPCR signaling complexes (Fig. 6b).

**Fig. 6.**
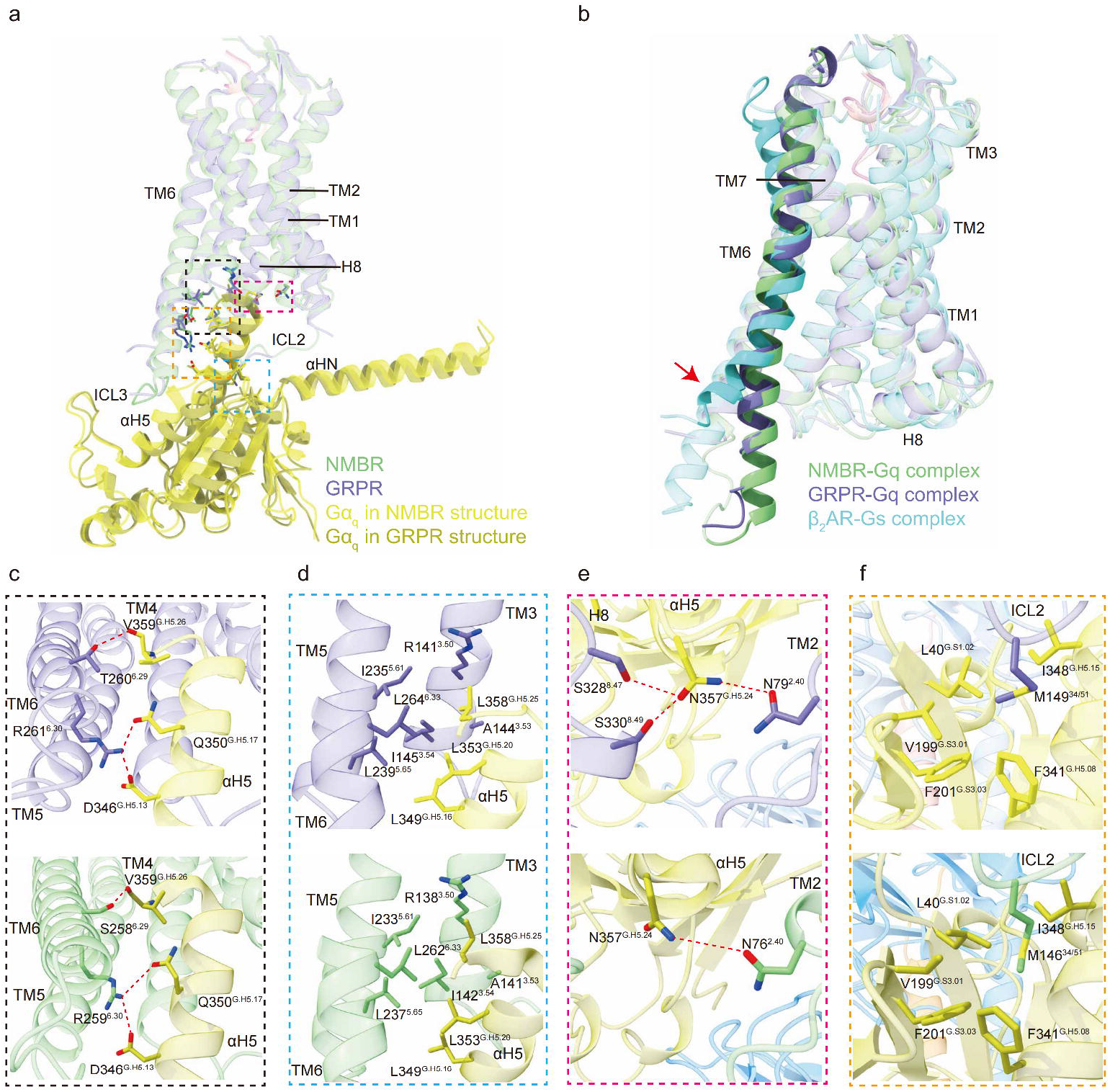
The G_q_ coupling mechanism of bombesin receptors. **a** Superposition of structures of the NMB30-NMBR-G_q_ and GRP (14-27)-GRPR-G_q_ complexes by aligning the receptors. The subunits Gβ and Gγ, together with the scFv16 in both structures are all omitted for clarity. **b** Structural superposition of two active bombesin receptors, active β_2_AR (PDB code: 3SN6) from the side view. Both bombesin receptors featured a less-pronounced outward swing movement of the intracellular tip of TM6 than that of the active G_s_-coupled β_2_AR, which is indicated by the red arrow. **c-f** Comparisons of detailed interactions of G_q_ and corresponding residues in NMBR and GRPR.

Aligned with the receptors, the two structures show a clear difference in the orientation and position of G protein relative to receptor, except for the very close conformation of the bound C-terminal alpha helix5 (αH5) of G_q_ (Supplementary information, Fig. S6d). When aligned with the G_q_ proteins, the two structures show a similar conformation of Gβγ, scFv16, and most regions of G_q_, except for the shift of the C-terminal αH5 and two receptors (Supplementary information, Fig. S6e, f). It is suggested that the αH5 can accommodate the different cytoplasmic TM cavities even in sub-family receptors, which plays a vital role in the G protein coupling. Indeed, the αH5 inserts into the cytoplasmic cavity and contributes to most of interactions with the receptor. The C-terminal carbonyl of V359^G.H5.26^ (superscription is CGN numbering system^25^) can form a hydrogen bond with the side chain of residue^6.29^ in both bombesin receptors (T260^6.29^ in NMBR and S258^6.29^ in GRPR), while the conserved L358^G.H5.25^ in the ‘wavy hook’ of G_q_ is blocked by the R^3.50^ to prevent further intrusion (Fig. 6c, d). At one side of the αH5, the side chains of conserved residues L349^G.H5.16^, L353^G.H5.20^, and L358^G.H5.25^ form a hydrophobic interaction network with nearby residues A^3.53^, I^3.54^, I^5.61^, L^5.65^ and L^6.33^ in both NMBR and GRPR structures (Fig. 6d). Adjacent to this conserved hydrophobic interaction network is the conserved residues R^6.30^ in both bombesin receptors, which forms electrostatic interaction with D346^G.H5.13^ and forms a hydrogen bond with Q350^G.H5.17^ from αH5 (Fig. 6c). In addition, at the top of the αH5 is the G_q_-family specific residue N^G.H5.24^ (G^G.H5.24^ in G_i/o_, M^G.H5.24^ in G_12/13_, and E^G.H5.24^ in G_s_ family) (Supplementary information, Fig. S8), which is reported to be important for receptor-G_q_ coupling^26^. In the NMBR structure, N^G.H5.24^ forms three hydrogen bonds with N79^2.40^ from TM2, S328^8.47^ and S330^8.49^ from TM7-H8 kink region. Similar interactions are also observed in the GRPR structure, and these specific receptor-G protein interactions may also contribute to selective coupling of bombesin receptors for the G_q_ subtype (Fig. 6e).

In addition, ICL2 and ICL3 are also reported to be important for G protein coupling. The conserved residue M^34.51^ (L^34.51^ in most G_q_-coupled receptors) in ICL2 are buried in a hydrophobic pocket formed by side chains of resides L40^G.S1.02^, V199^G.S3.01^, F201^G.S3.03^, F341^G.H5.08^, and I348^G.H5.15^ from G_q_ in both NMBR and GRPR structures (Fig. 6f). With the unambiguous density for the ICL2 and ICL3 regions in the GRPR structure, we can build all the residues confidently, which reveal extensive polar interactions between the ICL2 and the G_q_ in the GRPR structure, including Q149^34.54^ from ICL2 with R38^G.hns1.03^, S151^34.56^ from ICL2 with R37^G.hns1.02^ and Q34^G.HN.52^ (corresponding to the R34^G.HN.52^ in the WT G_q_ protein) (Supplementary information, Fig. S9a, b). The P245^5.73^ from ICL3 in in the GRPR structure forms aromatic packing with the side chains of Y325^G.S6.02^ and H327^G.S6.04^ (Supplementary information, Fig. S9c, d). In addition, there is a hydrogen bond interaction formed by the side chain of S72 from ICL1 with the side chain of D317 from Gβ in the NMBR structure (Supplementary information, Fig. S9e, f), yet there is no interaction between receptor and Gβ in the GRPR structure. In summary, we found the conserved and divergent features in G_q_ coupling to NMBR and GRPR.

### Activation mechanism of Bombesin Receptors

A structural comparison of NMBR and GRPR complexes to their closely related cholecystokinin A receptor (CCK_A_R) in the active (PDB 7EZM) and inactive states (PDB 7F8U) sheds light on the basis of bombesin receptor activation. Both NMBR and GRPR adopt fully active conformations similar to the active CCK_A_R (Fig. 7a). Compared with the inactive CCK_A_R structure, the active structures of NMBR and GRPR show a remarkable outward displacement of the cytoplasmic end of TM6, a hallmark of class A GPCR activation, along with an inward movement of the TM7 cytoplasmic end (Fig. 7a).

**Fig. 7.**
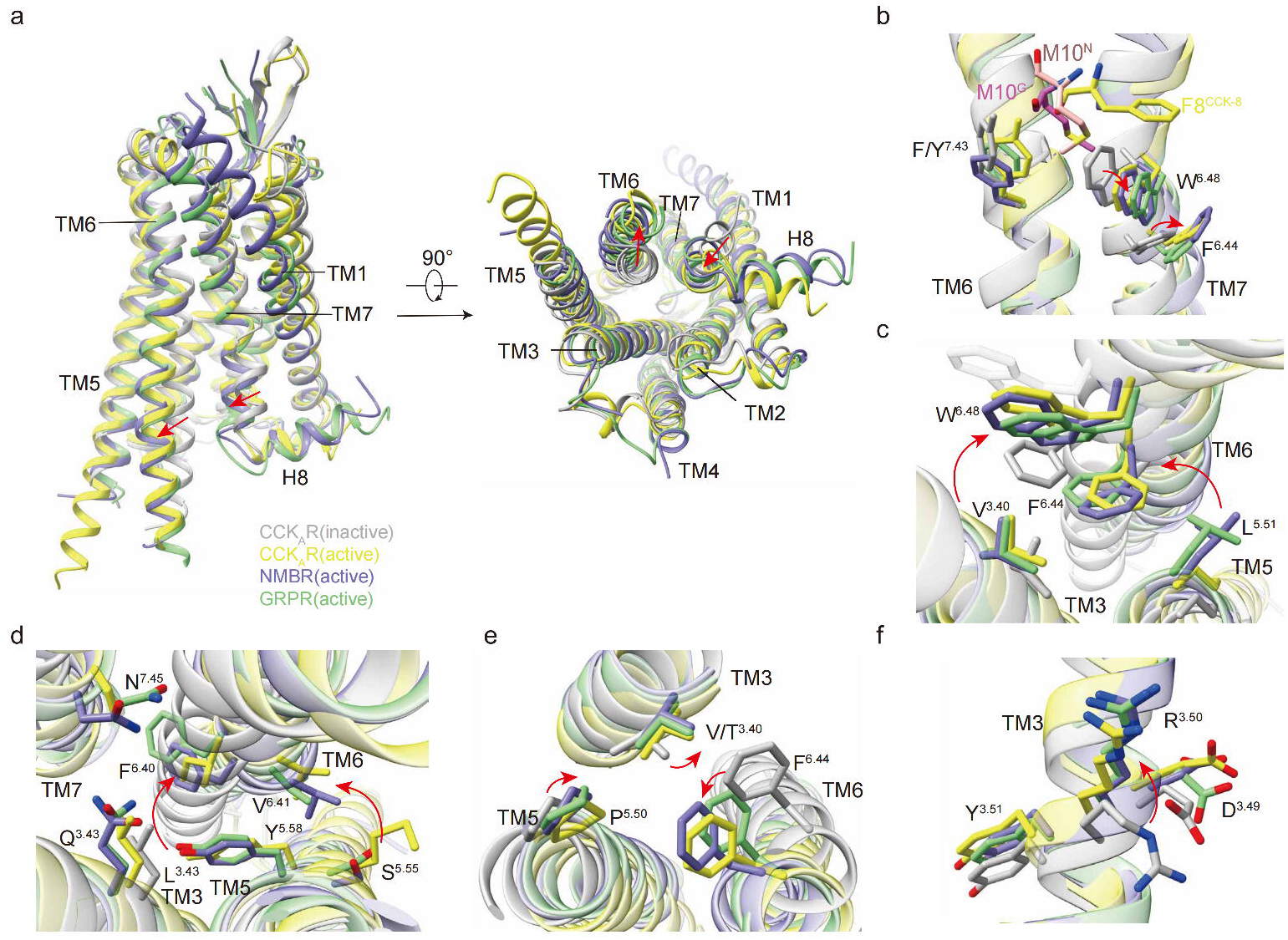
Molecular basis for activation of NMBR and GRPR. **a** Structural superposition of two active bombesin receptors, active CCK_A_R (PDB code: 7EZM) and inactive CCK_A_R (PDB code: 7F8U) from the side and cytoplasmic views. The movement directions of TM6 and TM7 in bombesin receptors relative to inactive CCK_A_R are highlighted as red arrows. CCK_A_R, Cholecystokinin A receptor. Inactive CCK_A_R, active CCK_A_R, NMBR, and GRPR are colored in gray, yellow, slate blue and green, respectively. **b-f** Conformational changes of the conserved ‘micro-switches’ upon receptor activation, including toggle switch (**b**), transmission switch consisting of L^5.51^ -F^6.44^ and V^3.40^-W^6.48^(**c**), conformational changes caused by L^3.43^Q constitutive mutation(**d**), PV(I)F (**e**), and DRY(**f**) motifs. The conformational changes of residue side chains are shown as red arrows upon receptor activation. The complex structures were aligned by the receptors.

Bombesin receptors could be activated by bombesin peptides with C termini both in amidated and non-amidated forms with different potencies. The amidated bombesin peptides presented nearly 100-fold higher potencies than the non-amidated forms for their corresponding receptors (Supplementary information, Fig. S7k, l and Table S3). Superimposing the NMBR and GRPR complexes reveals that the binding modes between the two bombesin peptides are very similar, suggesting that they may activate bombesin receptors through a common mechanism (Fig. 4c; Supplementary information, Fig. S6c, d), although NMB30 and GRP (14-27) present different binding selectivity. The side chains of M10^N^ and M10^G^ insert into a conserved hydrophobic crevice at the bottom of the peptide-binding pocket and trigger the downward rotameric switch of the toggle switch residue W^6.48^ in the conserved CWxP motif, facilitating the swing of F^6.44^ (Fig. 7b). Following the repacking of an intra-helical contact between residues at W^6.48^ and F^6.44^, the switching contacts of residues at V^3.40^ toward W^6.48^ and L^5.51^ toward F^6.44^ contract the TM3-5-6 interface (Fig. 7c), initializing the rotation of the cytoplasmic end of TM6.

As mentioned above, we introduced the L^3.43^Q mutation into both bombesin receptors to facilitate the formation of G-protein coupled complexes, which would loosen the hydrophobic lock (L^3.43^, F^6.40^, and V^6.41^ in bombesin receptors) and weaken TM3-TM6 contacts. Actually, both of the hydrophobic locks are broken in the two structures, instead F^6.40^, and V^6.41^ forms contacts with residues at Y^5.58^ and S^5.55^ (Fig. 7d), respectively. Besides, the introduced mutation L^3.43^Q could form a weak hydrogen bond with the side chain of N^7.45^ from the TM7 (Fig. 7d), which would promote the movement of TM7 toward TM3, and strengthen the packing of TM3-TM7. Finally, these switches further trigger the active-like conformational changes of ‘micro-switch’ residues (PV(I)F, DRY and NPxxL(Y) motifs) (Fig. 7e, f; Supplementary information, Table S4), leading to an agonism signal transduction to the cytoplasmic end of the receptor competent to bind to C terminus of Gα subunit.

The inward shift of TM7 to repack the TM3-TM7 interaction is a hallmark of class A GPCR activation, which is largely overlapped with that of other reported active conformation structures, such as ghrelin receptor (PDB 7F9Y^19^), B1R (PDB 7EIB^20^) and ET_B_ receptor (PDB 5GLH^27^). As shown in the M2R (PDB 4MQS^28^), Y^7.53^ in the conserved NPxxY motif forms a water-mediated hydrogen bond with Y^5.58^ to stabilizes this active conformation (Supplementary information, Fig. S10a). Nevertheless, these bombesin receptors both possess a substituted NPxxL motif with an infrequent L^7.53^, then such a hydrogen bond cannot be formed in the active conformation, which is also seen in the structure of ET_B_ receptor (have the same unusual NPxxL motif^27^) (Supplementary information, Fig. S10b), and the side chains of the equivalent L^7.53^ of these receptors display highly identical conformations. It is interesting that the inward shift of TM7 of bombesin receptors could be maintained in the absence of the hydrogen bond created by the conserved Y^7.53^ (Supplementary information, Fig. S10b). Thus, we believe that other residues around the cytoplasmic pocket would be responsible for alternative packing force to strengthen the packing of TM3-TM7. Overall, our study reveals the activation pathways of bombesin receptors and enriches the unique receptor-, ligand- and effector-specific activation pathways in the GPCR kingdoms.

## DISCUSSION

In this study, we present two cryo-EM structures of the active G_q_-coupled NMBR and GRPR bound to NMB30 and GRP (14-27), respectively. These structures reveal a unique squashed cavity in the extracellular side of these bombesin receptors, where three ECLs are folded to hold the resolved N-terminal segments of bombesin peptides. Intriguingly, the two bombesin peptides overlapped well in the orthosteric peptide-binding pocket, indicating a conserved mechanism of ligand recognition by subtype members of bombesin receptors. In addition, the binding modes of both NMB30 and GRP(14-27) are largely overlapped with other reported endogenous neuropeptides, including cholecystokinin (PDB 7EZM^29^), vasopressin (PDB 7DW9^30^), bradykinin (PDB 7F2O^20^) and orexin B (PDB 7L1U^31^) (Supplementary information, Fig. S10c). Superposition of these structures shows nearly identical conformations of C termini among different neuropeptides at the bottom of the pocket, while the extracellular sides undergo remarkable conformational shifts (Supplementary information, Fig. S10c). Considering the C-terminal sequences of most neuropeptides are related to receptor activation, this would partly explain the conserved neuropeptide receptor activation mechanism, of which is that the C terminus of neuropeptide is near the toggle switch residues to activate the receptor, while the N terminus is for selectivity and specificity to the receptor subtypes.

Both NMBR and GRPR could functionally constitute distinct but interconnected microcircuits to integrate and transmit itch signal from primary afferents to the brain^9^. The crucial roles of NMB/NMBR and GRP/GRPR signal pathways participated in itch perception and signal transduction established both NMBR and GRPR as tractable targets for therapeutic intervention. In addition, studies in human and animals suggest that both NMBR and GRPR are involved in a broad range of physiological and pathophysiological processes, including contraction of smooth muscle, food intake, release of some gastrointestinal hormones and inflammation. Furthermore, the NMBR and GRPR are two of the G protein-coupled receptors that are most frequently overexpressed by various human neoplasms including cancers of the lung, pancreas, prostate, CNS, head/neck, breast, and various neuroendocrine tumors^32,33^. As such both NMBR and GRPR are recognized as neoplasm biomarkers. Recently there has been increased interest in developing high affinity ligands that are metabolically stable and can be linked to various effectors, including radiolabel ligands for imaging bombesin receptors-overexpressed tumors and cytotoxic compounds for antitumor treatment. Our structures depict the detailed interactions between the endogenous ligands and bombesin receptors. With the in-depth knowledge of ligand binding and selectivity of bombesin receptors, new opportunities will arise to design potent and efficacious modulators of bombesin receptors for the treatment of pruritus and carcinoma.

## AUTHOR CONTRIBUTIONS

C.L. screened the expression constructs, purified the bombesin receptor complex proteins, participated in cryo-EM grid inspection, data collection, designed the mutations and executed the functional studies. Y.X. prepared the cryo-EM grids, collected cryo-EM images with the help of Q.Y., K.W. and W.H., and performed density map calculations and participated in the model building and refined the final models. Y.X. and H.L. participated in figure and manuscript preparation. H.E.X. and W.Y. conceived and supervised the project. W.Y. and C.L. prepared the figures and drafted the manuscript. H.E.X. and W.Y. wrote the manuscript with input from all authors.

## ACKNOWLEDGMENTS

The cryo-EM data of the bombesin receptor complex proteins were collected at the Shanghai Advanced Electron Microscope Center, Shanghai Institute of Material Medica. We thank all the staff (Q.Y., K.W. and W.H.) at the cryo-EM facility for their technical support. This work was partially supported by Ministry of Science and Technology (China) grants (2018YFA0507002 to H.E.X.); Shanghai Municipal Science and Technology Major Project (2019SHZDZX02 to H.E.X.); Shanghai Municipal Science and Technology Major Project (H.E.X.); CAS Strategic Priority Research Program (XDB37030103 to H.E.X.), National Natural Science Foundation of China (32130022 to H.E.X.); the Youth Innovation Promotion Association of CAS (2021278 to W.Y.); National Natural Science Foundation of China (32171189 to W.Y.); National Science Fund for Excellent Young Scholars (82122067 to W.Y.); Key tasks of the Lingang Laboratory (LG202103-03-05 to W.Y.); Key tasks of Lingang Laboratory (LG202101-01-03 to Y. X.); China Postdoctoral Science Foundation (2021M703341 to H.C.); National Natural Science Foundation (31770796 to Y.J.); and National Science and Technology Major Project (2018ZX09711002 to Y.J.). In addition, this work was partially supported by High-level new R&D institute (2019B090904008), and High-level Innovative Research Institute (2021B0909050003) from Department of Science and Technology of Guangdong Province, and the author W.Y. also gratefully acknowledges the support of Sanofi Scholarship Program.

## Competing interests

The authors declare no competing interests.

## Data Resources

Materials are available from the corresponding authors upon reasonable request. Density maps and structure coordinates have been deposited in the Electron Microscopy Data Bank (EMDB) and the Protein Data Bank (PDB) with accession codes EMD-34413 and 8H0P for NMB30-NMBR-Gq complex; and EMD-34414 and 8H0Q for GRP (14-27)-GRPR-Gq complex. Source data are provided with this paper.

## Notes

### Competing Interest Statement

The authors have declared no competing interest.

