## supplementary information-methods for "Molecular Recognition of Itching Neuropeptides by Bombesin Receptors"

**Constructs of NMBR, GRPR, and G proteins**

Human NMBR (residues 1-361) and GRPR (residues 1-359) were cloned into pFastBac with an N-terminal haemagglutinin(HA) signal peptide followed by thermostabilized apocytochrome b_562_RIL (BRIL)^1^ as well as LgBiT at the C-terminal using homologous recombination (CloneExpress One Step Cloning Kit, Vazyme). After LgBiT, a TEV protease cleavage site and a double-MBP tag were added to both NMBR and GRPR constructs used for better expression and purification, and the mutation L134^3.43^Q of NMBR and L131^3.43^Q of GRPR were generated by site-directed mutagenesis to facilitate receptor selectivity to Gαq and complex stability^2^. The engineered Gα_q_ construct was generated based on mini-Gs/q71^3^, which carries two dominant-negative mutations (corresponding to G203A and A326S)^4^ to decrease the affinity of nucleotide-binding and facilitate the stability of Gαβγ complex. The N terminal 1–18 amino acids and the α-helical domain of the mini-G_s/q_71 were substituted by the corresponding sequences of the human Gαi1 contributing to binding to the antibody fragments scFv16^5, 6^. Gβ1 was connected with a C-terminal SmBiT by a 15 residues linker. The engineered Gα_q_, Gβ1-SmBiT and Gγ2 were cloned into pFastBac vectors independently (Invitrogen).

**Expression and purification of NMBR/GRPR–G_q_ complex.**

High Five cells (Expression systems) were cultured in ESF921 serum-free medium (Expression Systems) and infected with viruses of the receptor (NMBR or GRPR), Gα_q_, Gβ1, Gγ2, and scFv16^7^ in the ratio of 1:1:1:1:1 for 48 h at 27 ˚C using Bac-to-Bac baculovirus system. The cell pellets were lysed by dounce homogenization in a buffer containing 20 mM HEPES pH 7.4, 100 mM NaCl, 10 mM MgCl_2_, 5 mM CaCl_2_, 0.1 mM TCEP (Sigma-Aldrich), 10% glycerol, and EDTA-free protease inhibitor cocktail (TargetMol). The supernatant was isolated by centrifugation at 65,000 × g for 40 min to collect the membranes. The washed membranes were re-suspended in 20 mM HEPES pH 7.4, 100 mM NaCl, 10 mM MgCl_2_, 5 mM CaCl_2_, 10% glycerol, 10 μM peptide (NMB30 for NMBR and GRP (14-27) for GRPR, respectively, Synpeptide), 25 mU/mL apyrase (Sigma-Aldrich), 0.1 mM TCEP and EDTA-free protease inhibitor cocktail, and incubated at room temperature for 1.5 h. After incubation, 0.5% (w/v) n-dodecyl-β-d-maltopyranoside (DDM, Anatrace) and 0.1% (w/v) cholesteryl hemisuccinate TRIS salt (CHS, Anatrace) was used for solubilization at 4˚C for 2.5 h. The supernatant was collected by centrifugation at 65,000 × g for 40 min and then incubated with dextrin resin (Dextrin Beads 6FF, Smart Life Sciences) at 4˚C for 4 h. The resin was collected by centrifugation at 500 × g for 5 min, loaded onto a gravity flow column and washed with 10 column volumes of buffer containing 20 mM HEPES pH 7.4, 100 mM NaCl, 10 mM MgCl_2_, 5 mM CaCl_2_, 10% glycerol, 0.1 mM TCEP, 5 μM peptide, 0.05% (w/v) DDM and 0.01% (w/v) CHS. The detergent of washing buffer was then displaced by 0.1% (w/v) lauryl maltose neopentylglycol (LMNG, Anatrace) and 0.02% (w/v) CHS for 10 column volumes washing, followed by 0.03% (w/v) LMNG, 0.01% (w/v) glyco-diosgenin (GDN, Anatrace) and 0.008% (w/v) CHS for 20 column volumes washing. The protein was then treated with His-tagged TEV protease on column and further incubated at 4 ˚C for 8 h. The elution was concentrated with an Amicon Ultra Centrifugal Filter (MWCO 100 kDa) and injected onto a Superdex 200 increase 10/300 GL column (GE Healthcare) with running buffer 20 mM HEPES pH 7.4, 100 mM NaCl, 2 mM MgCl_2_, 0.1 mM TCEP, 5 μM peptide, 0.00075%(w/v) LMNG, 0.00025% (w/v) GDN and 0.0002% (w/v) CHS. The fractions of monomeric protein complex for NMBR and GRPR were collected and evaluated by SDS-PAGE (Supplementary information, Fig. S2), and then concentrated by 30–50-fold for cryo-electron microscopy experiments.

**Cryo-EM data collection**

Cryo-EM grids were prepared with the Vitrobot Mark IV plunger (FEI) set to 4 ˚C and 100% humidity. Three-microliter of the NMB30-NMBR-G_q_ complex was applied to the glow discharged Au R1.2/1.3 holey carbon grids. The sample was incubated for 5 s on the grids before blotting for 3 s (double-sided, blot force 2) and flash-frozen in liquid ethane immediately. The same condition was used for the sample GRP (14-27)-GRPR-G_q_ complex.

For NMB30-NMBR-G_q_ complex dataset, 4,858 movies were collected on a Titan Krios equipped with a Gatan K3 direct electron detection device at 300 kV with a magnification of 81,000, corresponding to a pixel size 1.04 Å. Image acquisition was performed with EPU Software (FEI Eindhoven, Netherlands). We collected a total of 36 frames accumulating to a total dose of 50 e^-^ Å^-2^ over 2.5 s exposure.

For GRP (14-27)-GRPR-G_q_ complex dataset, 9,002 movies were collected on a Titan Krios equipped with a Falcon4 direct electron detection device at 300 kV with a magnification of 96,000, corresponding to a pixel size 0.8 Å. Image acquisition was performed with EPU Software (FEI Eindhoven, Netherlands). We collected a total dose of 50 e^-^ Å^-2^ over 2.5 s exposure on each EER format movie^8^. Each movie was divided into 36 frames during motion correction.

**Cryo-EM image processing**

MotionCor2 was used to perform the frame-based motion-correction algorithm to generate drift-corrected micrograph for further processing and CTFFIND4 provided the estimation of the contrast transfer function (CTF) parameters^9, 10^.

For NMB30-NMBR-G_q_ complex dataset, 480 aligned micrographs were deleted because of contaminations or bad ice quality. After selection, approximately 800 particles were manually picked and 2D classes were calculated and used as references for automatic picking. All subsequent steps of particle picking, extraction, classification and post processing of refined models were performed with Relion3.0^11^. A total of 3,034,736 particles were extracted from the cryo-EM micrographs and followed by reference-free two-dimensional (2D) classification, yielding 619,210 particles after clearance. Mask three-dimensional (3D) classification on the receptor part was used to separate out 355,509 particles that resulted to a clearer density of NMBR. We refined this portion of particles, which led to a structure at 3.52 Å global resolution. After CTF refinement, Bayesian polishing, and postprocessing with DeepEMhancer^12^, then the particles were reconstituted to a 3.15 Å structure (Supplementary information, Fig. S3).

For GRP (14-27)-GRPR-G_q_ complex dataset, 762 aligned micrographs were deleted because of contaminations or bad ice quality. After selection, NMBR was used as 3D reference for automatic picking. All subsequent steps of particle picking, extraction, classification and post processing of refined models were performed with Relion3.0^11^. A total of 3,365,839 particles were extracted from the cryo-EM micrographs and followed by reference-free two-dimensional (2D) classification, yielding 577,108 particles after clearance. Mask three-dimensional (3D) classification on the receptor part was used to separate out 301,192 particles that resulted to a clearer density of GRPR. The second round of 3D classification was performed without mask and separated out 55,286 particles. We refined the remained particles, which led to a structure at 3.72 Å global resolution. After the postprocessing with DeepEMhancer^12^, then the particles were reconstituted to a 3.3 Å structure (Supplementary information, Fig. S4).

**Model building**

NMBR and GRPR structures predicted from Alphafold2 were used as the starting reference models for receptors building^13^. Structures of Gα_q_, Gβ, Gγ and the scFv16 were derived from PDB entry 7WKD (unpublished) were rigid body fit into the density. All models were fitted into the EM density map using UCSF Chimera^14^ followed by iterative rounds of manual adjustment and automated rebuilding in COOT^15^ and PHENIX^16^, respectively. The model was finalized by rebuilding in ISOLDE^17^ followed by refinement in PHENIX with torsion-angle restraints to the input model. The final model statistics were validated using Comprehensive validation (cryo-EM) in PHENIX^16^ and provided in the supplementary information (Supplementary information, Table S1). All structural figures were prepared using Chimera^14^, Chimera X^18^, and PyMOL (Schrödinger, LLC.).

**Function essay**

AD293 cells (Agilent) were cultured in DMEM/high Glucose medium (GE healthcare) supplemented with 10% (v/v) fetal bovine serum (FBS, Gemini) and 1% penicillin/streptomycin and maintained at 37°C in 5% CO_2_ incubator. Inositol phosphate 1 (IP1) production was measured using the IP-One HTRF kit (Cisbio, 621PAPEJ)^19^. Briefly, cells were seeded onto 12-well cell culture plates for 16 h before transfection. The cells were then transiently with different NMBR or GRPR constructs using FuGENE HD transfection reagent. After 24 h, cells were harvested and resuspended in IP1 stimulation buffer at a density of 7 × 10^5^ cells/mL. Cells were then plated onto 384-well assay plates at 4900 cells/7 μL/well. Another 7 μL IP1 Stimulation Buffer 2 containing ligand was added to the cells, and the incubation lasted for 1 h at 37 °C. Intracellular IP1 measurement was carried with the IP-One HTRF kit and EnVision multiplate reader according to the manufacturer’s instructions. The HTRF ratio was converted to a response (%) using the following formula: response (%) = ratio of sample/WT×100. Data presented are mean±S.E.M. of at least three independent experiments.

**Cell-surface expression assay**

Cell-surface expression for each NMBR and GRPR mutant was monitored by a fluorescence-activated cell sorting (FACS) assay. The mutants were cloned into pcDNA6.0 vector (Invitrogen) with a N-terminal FLAG tag. The cell seeding and transfection follow the same method as function assay. After 24h of transfection, cells were washed once with PBS and digested with 0.2% (w/v) EDTA in PBS. Thereafter, the expressed cells were incubated with Monoclonal anti-FLAG M2-FITC (Sigma-Aldrich) at a dilution of 1:100 for 15 min at 4 °C, and then a 9-fold excess of PBS was added to cells. After cells were resuspended, fluorescence intensity was quantified in a BD Accuri C6 flow cytometer system (BD Biosciences) at excitation 488 nm and emission 519 nm. The FACS data were analyzed by BD Accuri C6 software 1.0.264.21 and data were normalized to WT.

**Figure Legends for supplementary figures**

**Fig. S1** **Sequence alignment of NMBR, GRPR and BRS3. Secondary structure elements are annotated underneath the sequences based on the structure of the NMB30-NMBR-G_q_ complex.**

**Fig. S2** **Purification and characterization of NMB30-NMBR–G_q_ complex and GRP (14-27)-GRPR-G_q_ complex.**

**a** Schematic diagram of the protein engineering of NMBR, engineered Gα_q_, Gβ, Gγ and scFv16 used in this study.

**b** Gel filtration (Superdex 200 Increase 10/300 column, GE Healthcare) profile of the NMB30-NMBR–G_q_ complex. The black arrow indicates the sharp peak for NMB30-NMBR–G_q_ complex.

**c** Coomassie-stained SDS-PAGE analysis of the purified NMB30-NMBR–G_q_ complex, showing balanced ratios for each subunit.

**d** Schematic diagram of the protein engineering of NMBR, engineered Gα_q_, Gβ, Gγ and scFv16 used in this study.

**e** Gel filtration (Superdex 200 Increase 10/300 column, GE Healthcare) profile of the GRP (14-27)-GRPR-G_q_ complex. The black arrow indicates the sharp peak for GRP (14-27)-GRPR-G_q_ complex.

**f** Coomassie-stained SDS-PAGE analysis of the purified GRP (14-27)-GRPR-G_q_ complex, showing balanced ratios for certain subunits, except for the Gγ subunit.

**Fig. S3** **Cryo-EM data processing of NMB30-NMBR-G_q_ complex.**

**a** Flowchart of computational sorting of cryo-EM data.

**b** A representative cryo-EM micrograph of NMB30-NMBR–Gq complex with 50 nm scale bar included as a size reference.

**c** Twelve representative reference-free 2D cryo-EM class averages. Scale bar, 5 nm.

**d** ‘Gold-standard’ Fourier shell correlation curve of the reconstruction. The resolution was reported at 3.15 Å using the Fourier shell cutoff at 0.143.

**e** Local resolution map of NMB30-NMBR–Gq complex.

**Fig. S4** **Cryo-EM data processing of GRP (14-27)-GRPR-G_q_ complex.**

**a** Flowchart of computational sorting of cryo-EM data.

**b** A representative cryo-EM micrograph of GRP (14-27)-GRPR-G_q_ complex with 50 nm scale bar included as a size reference.

**c** Twelve representative reference-free 2D cryo-EM class averages. Scale bar, 5 nm.

**d** Gold-standard’ Fourier shell correlation curve of the reconstruction. The resolution was reported at 3.3 Å using the Fourier shell cutoff at 0.143.

**e** Local resolution map of GRP (14-27)-GRPR-G_q_ complex.

**Fig. S5** **Cryo-EM density maps of TM1-7 in receptors, αH5 and αHN in G_q_ proteins， and ligands in NMBR structure (a) and GRPR structure (b).**

**Fig. S6** **Structural alignment of the structures of the NMB30-NMBR-G_q_ complex and GRP (14-27)-GRPR-G_q_ complex.**

**a** Structural alignment of overall structures of NMB30-NMBR-G_q_ and GRP (14-27)-GRPR-G_q_ complex.

**b** Structural alignment of the structures of NMBR and GRPR by aligning the receptors.

**c** Superposition of structures of the NMB30-NMBR-G_q_ and GRP (14-27)-GRPR-G_q_ complexes by aligning the receptors in a different view related to Figure 6A. The subunits Gβ and Gγ, together with the scFv16 in both structures are all omitted for clarity.

**d** Superposition of structures of the NMB30-NMBR-Gq and GRP (14-27)-GRPR-Gq complexes by aligning the receptors, showing the different orientations and positions of G proteins.

**e,f** Superposition of structures of the NMB30-NMBR-Gq and GRP (14-27)-GRPR-Gq complexes by aligning the G_q_ proteins, showing the different orientations of the two receptors (**e**) and the shift of the C-terminal αH5 (**f**).

**Fig. S7** **IP1 response curves of NMBR and GRPR.**

Effects of NMBR mutations (**a-e**) or GRPR mutations (**f-j**) on NMB30 or GRP (14-27)-induced IP production. Effects of amidated and non-amidated forms of NMB30 or GRP (14-27) on receptor activation (**k, l**). Data are presented as mean ± S.E.M. of at least three independent experiments. Source data are available online.

**Fig. S8** **Sequence alignment of Gα_q_, engineered Gα_q_, Gα_11_, Gα_s_, Gα_i1_, Gα_o_, Gα_12_, and Gα_13_.**

**Fig. S9** **Detailed interactions of ICLs from NMBR and GRPR wtih G_q_ proteins.**

**a, b** Cryo-EM map and interaction diagram of ICL2 in GRPR and αN helix in Gα_q_.

**c, d** Cryo-EM maps and interaction diagram of ICL3 in GRPR and Gα_q_.

**e, f** Cryo-EM maps and interaction diagram of ICL1 in NMBR and Gβ.

**Fig. S10 Molecular basis for activation of NMBR and GRPR and structure comparison of NMB30 and GRP (14-27) with other neuropeptides solved.**

**a, b** The structural features of the conserved NPXXY motifs in GHSR (PDB code: 7F9Y), B1R (PDB code: 7EIB) and M2R (PDB code: 4MQS) (**a**), and the substituted NPXXL motifs with the infrequent L^7.53^ in ET_B_ (PDB code: 5GLH), NMBR and GRPR (H). The putative water molecule in conserved NPXXY motif is shown (**b**).

**c** Structure comparison of NMB30 and GRP (14-27) with other neuropeptides solved. The neuropeptides are shown as a cartoon. The shift of the extracellular part of neuropeptides is highlighted as a red arrow. The neuropeptide colors are indicated as the labeled receptors: vasopressin (PDB code: 7DW9), cyan; CCK-8 (PDB code: 7EZM), green; orexin B (PDB code: 7L1U), orange; bradykinin (PDB code: 7F2O), red.

**Table S1** **Cryo-EM data collection, model refinement and validation statistics of the NMB30-NMBR-G_q_ complex and GRP (14-27)-GRPR-G_q_ complex.**

**Table S2** **Amino acid sequences of Bn-related peptides used or mentioned in this study.**

**Table S3** **Ligand binding affinities and expression levels of WT and mutated NMBR and GRPR. The wild type (WT) and mutants of NMBR and GRPR discussed in this manuscript were individually analyzed. The affinities are derived from at least 3 independent experiments using IP1 function assay. The expression level of mutant NMBR and GRPR were normalized to wild-type NMBR and GRPR as 100%, respectively. Each data point represents mean ± standard error of the mean (S.E.M.). All data were analyzed by two-sided Student’s t test. *P<0.05, **P<0.01, ***P<0.001 vs. WT. Source data are available online. Definitions: NA – not applicable; NT, not tested.**

**Table S4** **Sequence alignment of the key residues in sodium site, DRY motif, PV(I)F motif, toggle switch and NPxxP motif, as well as residues involved in disulfide bond formation in bombesin receptors.**
