## supplementary figures and tables for "Molecular Recognition of Itching Neuropeptides by Bombesin Receptors"

### Fig. S1

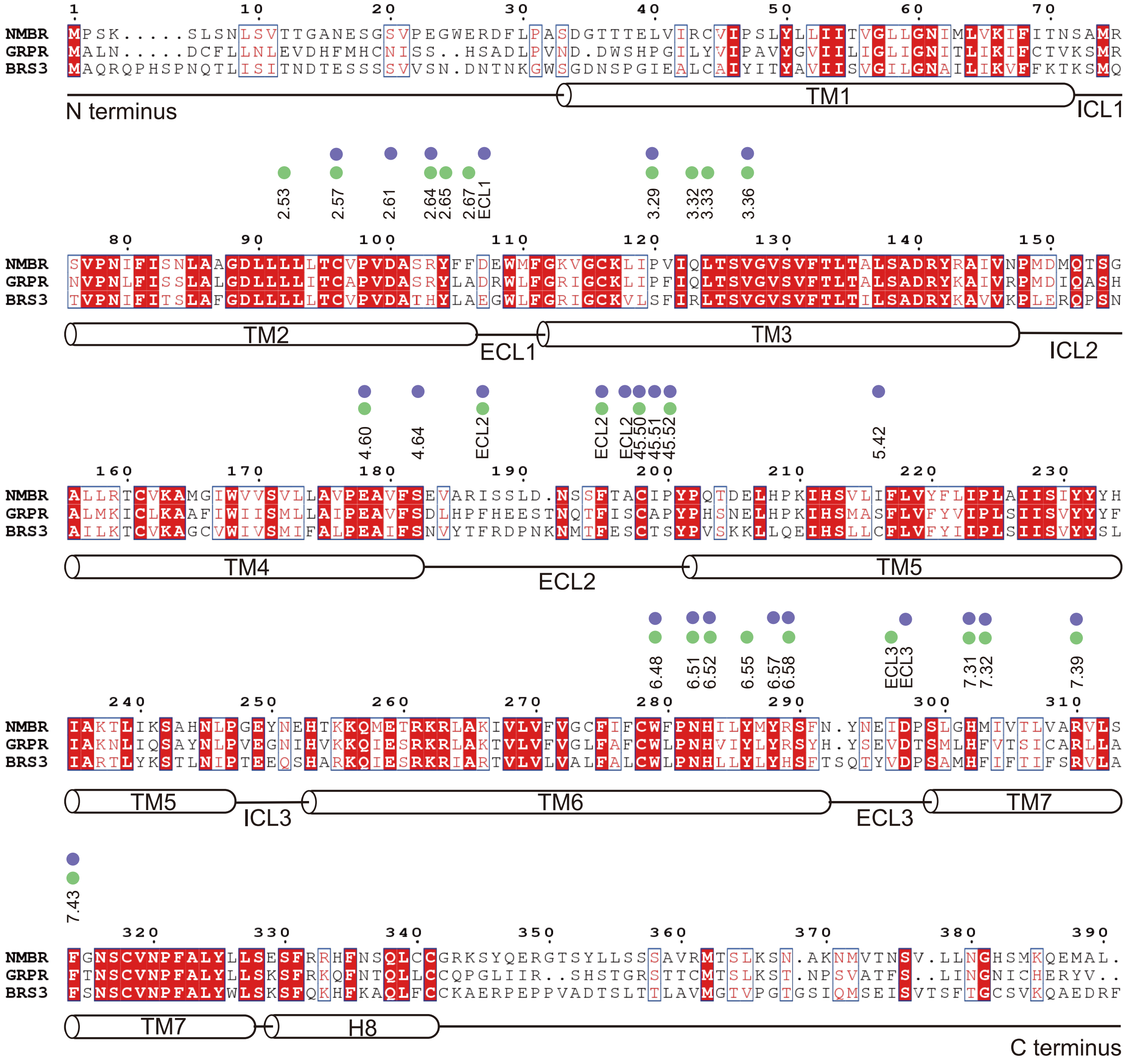

NMBR MPSK.....SLSNLSVTTGANESGSVPEGWERDFLPASDGTTTELVIRCVIPSLYLLIITVGLLGNIMLVKIFITNSAMR
GRPR MALN.....DCFLLNLEVDHFMHCNISS..HSADLPVND.DWSHPGILYVIPAVYGVIILIGLIGNITLIKIFCTVKSMR
BRS3 MAQRQPHSPNQTLISITNDTESSSSVVSN.DNTNKGWSGDNSPGIEALCAIYITYAVIISVGILGNAILIKVFFKTKSMQ
N terminus
TM1
ICL1
2.53 2.57 2.61 2.64 2.65 2.67 ECL1 3.29 3.32 3.33 3.36
NMBR SVPNIFISNLAAGDLLLLLTCVPVDASRYFFDEWMFGKVGCKLIPVIQLTSVGVSVFTLTALSADRYRAIVNPMDMQTSG
GRPR NVPNLFISSLALGDLLLLITCAPVDASRYLADRWLFGRIGCKLIPFIQLTSVGVSVFTLTALSADRYKAIVRPMDIQASH
BRS3 TVPNIFITSLAFGDLLLLLTCVPVDATHYLAEGWLFGRIGCKVLSFIRLTSVGVSVFTLTILSADRYKAVVKPLERQPSN
TM2
ECL1
TM3
ICL2
4.60 4.64 ECL2 ECL2 ECL2 45.50 45.51 45.52 5.42
NMBR ALLRTCVKAMGIWVVSVLLAVPEAVFSEVARISSLD.NSSFTACIPYPQTDELHPKIHSVLIFLVYFLIPLAIISIYYYH
GRPR ALMKICLKAAFIWIISMLLAIPEAVFSDLHPFHEESTNQTFISCAPYPHSNELHPKIHSMASFLVFYVIPLSIISVYYYF
BRS3 AILKTCVKAGCVWIVSMIFALPEAIFSNVYTFRDPNKNMTFESCTSYPVSKKLLQEIHSLLCFLVFYIIPLSIISVYYSL
TM4
ECL2
TM5
6.48 6.51 6.52 6.55 6.57 6.58 ECL3 ECL3 7.31 7.32 7.39
NMBR IAKTLIKSAHNLPGEYNEHTKKQMETRKRLAKIVLVFVGCFIFCWFPNHILYMYRSFN.YNEIDPSLGHMIVTLVARVLS
GRPR IAKNLIQSAYNLPVEGNIHVKKQIESRKRLAKTVLVFVGLFAFCWLPNHVIYLYRSYH.YSEVDTSMLHFVTSICARLLA
BRS3 IARTLYKSTLNIPTEEQSHARKQIESRKRIARTVLVLVALFALCWLPNHLLYLYHSFTSQTYVDPSAMHFIFTIIFSRVLA
TM5
ICL3
TM6
ECL3
TM7
7.43
NMBR FGNSCVNPFALYLLSESFRRHFNSQLCCGRKSYQERGTSYLLSSSAVRMTSLKSN.AKNMVTNSV.LLNGHSMKQEMAL.
GRPR FTNSCVNPFALYLLSKSFRKQFNTQLLCCQPGLIIR..SHSTGRSTTCMTSLKST.NPSVATFS..LINGNICHERYV..
BRS3 FSNSCVNPFALYWLSKSFQKHFKAQLFCCKAERPEPPVADTSLTTLAVMGTVPGTGSIQMSEISVTSFTGCSVKQAEDRF
TM7
H8
C terminus

Fig. S2

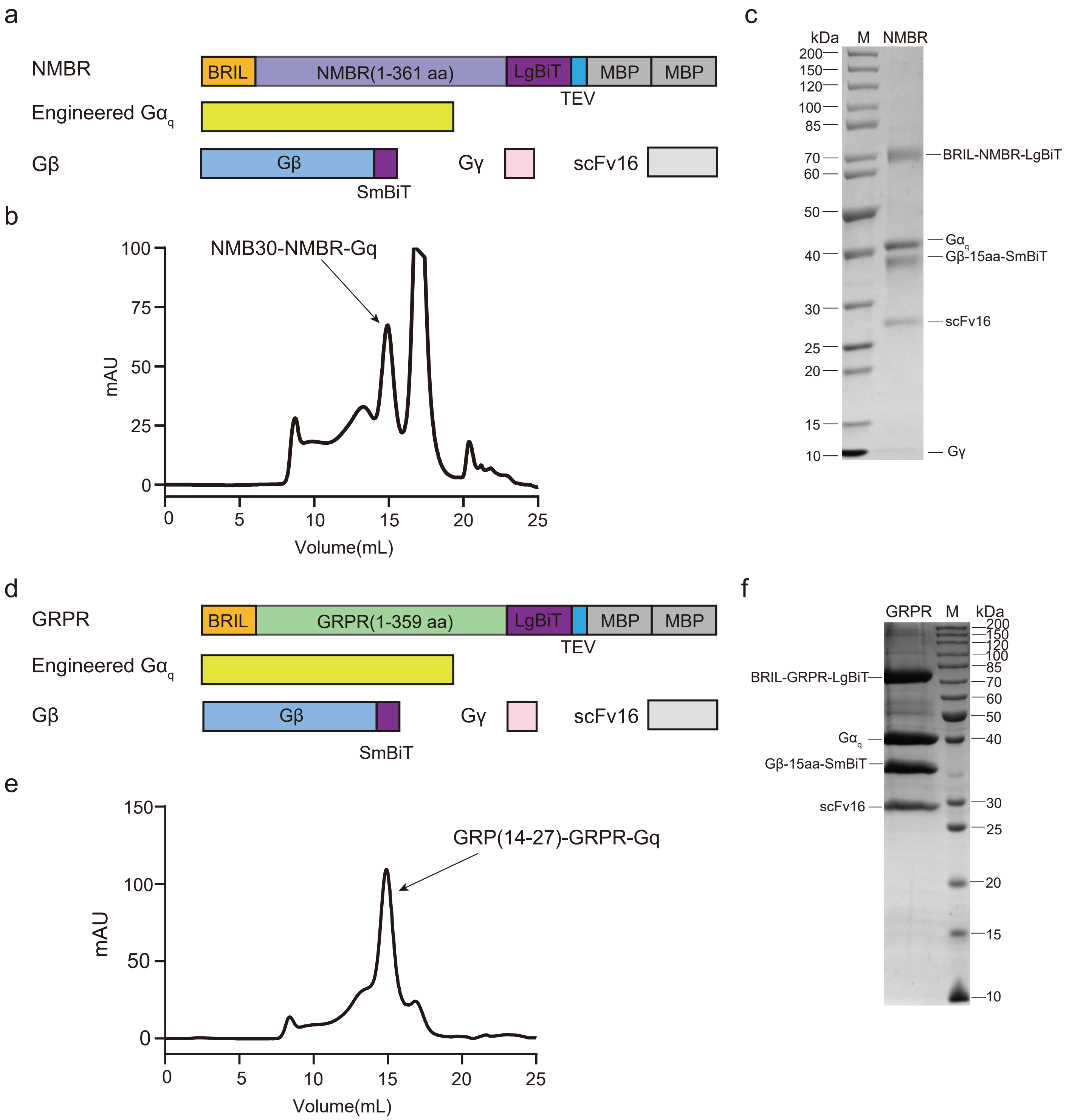
a
NMBR
BRIL
NMBR(1-361 aa)
LgBiT
MBP
MBP
TEV
Engineered Gα$_q$
Gβ
Gβ
SmBiT
Gγ
scFv16
b
NMB30-NMBR-Gq
mAU
100
75
50
25
0
0
5
10
15
20
25
Volume(mL)
c
kDa
M
NMBR
200
150
120
100
85
70
60
50
40
30
25
20
15
10
BRIL-NMBR-LgBiT
Gα$_q$
Gβ-15aa-SmBiT
scFv16
Gγ
d
GRPR
BRIL
GRPR(1-359 aa)
LgBiT
MBP
MBP
TEV
Engineered Gα$_q$
Gβ
Gβ
SmBiT
Gγ
scFv16
e
GRP(14-27)-GRPR-Gq
mAU
150
100
50
0
0
5
10
15
20
25
Volume(mL)
f
GRPR
M
kDa
200
150
120
100
85
70
60
50
40
30
25
20
15
10
BRIL-GRPR-LgBiT
Gα$_q$
Gβ-15aa-SmBiT
scFv16

Fig. S3

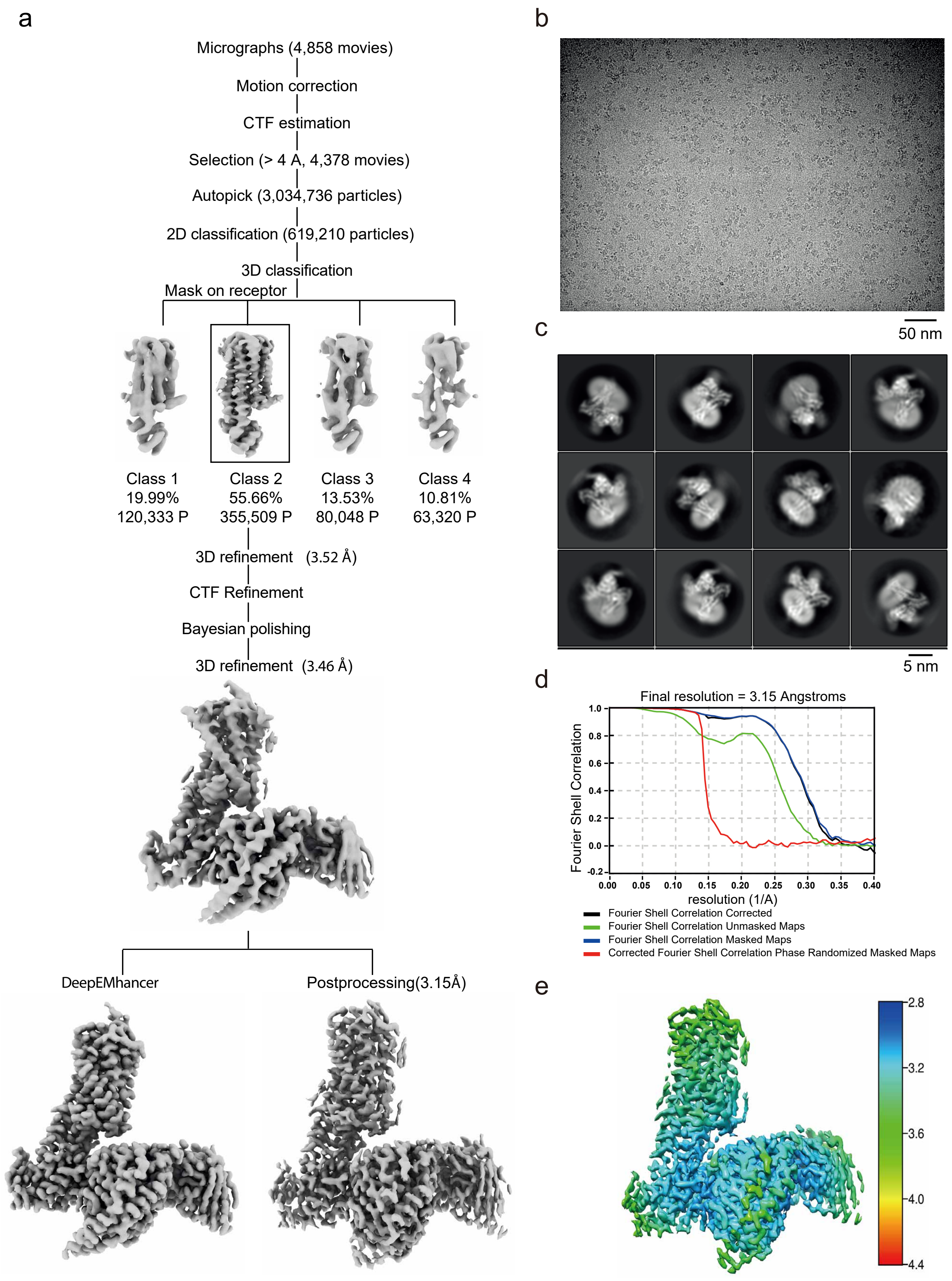

a
Micrographs (4,858 movies)
Motion correction
CTF estimation
Selection (> 4 A, 4,378 movies)
Autopick (3,034,736 particles)
2D classification (619,210 particles)
3D classification
Mask on receptor
Class 1
19.99%
120,333 P
Class 2
55.66%
355,509 P
Class 3
13.53%
80,048 P
Class 4
10.81%
63,320 P
3D refinement (3.52 Å)
CTF Refinement
Bayesian polishing
3D refinement (3.46 Å)
DeepEMhancer
Postprocessing(3.15Å)
b
50 nm
c
5 nm
d
Final resolution = 3.15 Angstroms
Fourier Shell Correlation
resolution (1/A)
Fourier Shell Correlation Corrected
Fourier Shell Correlation Unmasked Maps
Fourier Shell Correlation Masked Maps
Corrected Fourier Shell Correlation Phase Randomized Masked Maps
e
2.8
3.2
3.6
4.0
4.4

Fig. S4

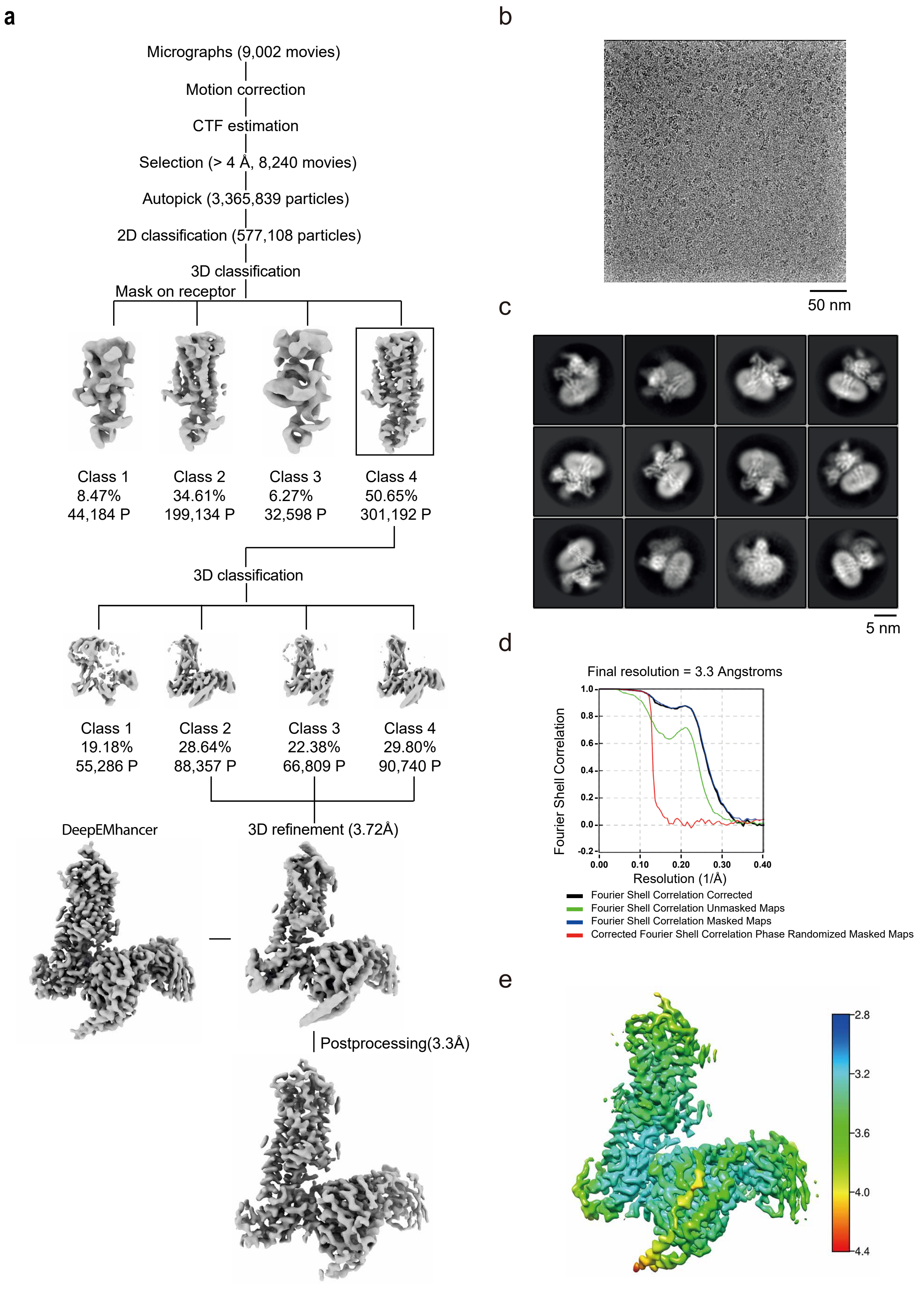
a
Micrographs (9,002 movies)
Motion correction
CTF estimation
Selection (> 4 Å, 8,240 movies)
Autopick (3,365,839 particles)
2D classification (577,108 particles)
3D classification
Mask on receptor
Class 1
8.47%
44,184 P
Class 2
34.61%
199,134 P
Class 3
6.27%
32,598 P
Class 4
50.65%
301,192 P
3D classification
Class 1
19.18%
55,286 P
Class 2
28.64%
88,357 P
Class 3
22.38%
66,809 P
Class 4
29.80%
90,740 P
DeepEMhancer
3D refinement (3.72Å)
Postprocessing(3.3Å)
b
50 nm
c
5 nm
d
Final resolution = 3.3 Angstroms
Fourier Shell Correlation
Resolution (1/Å)
1.0
0.8
0.6
0.4
0.2
0.0
-0.2
0.00
0.10
0.20
0.30
0.40
Fourier Shell Correlation Corrected
Fourier Shell Correlation Unmasked Maps
Fourier Shell Correlation Masked Maps
Corrected Fourier Shell Correlation Phase Randomized Masked Maps
e
2.8
3.2
3.6
4.0
4.4

Fig. S5

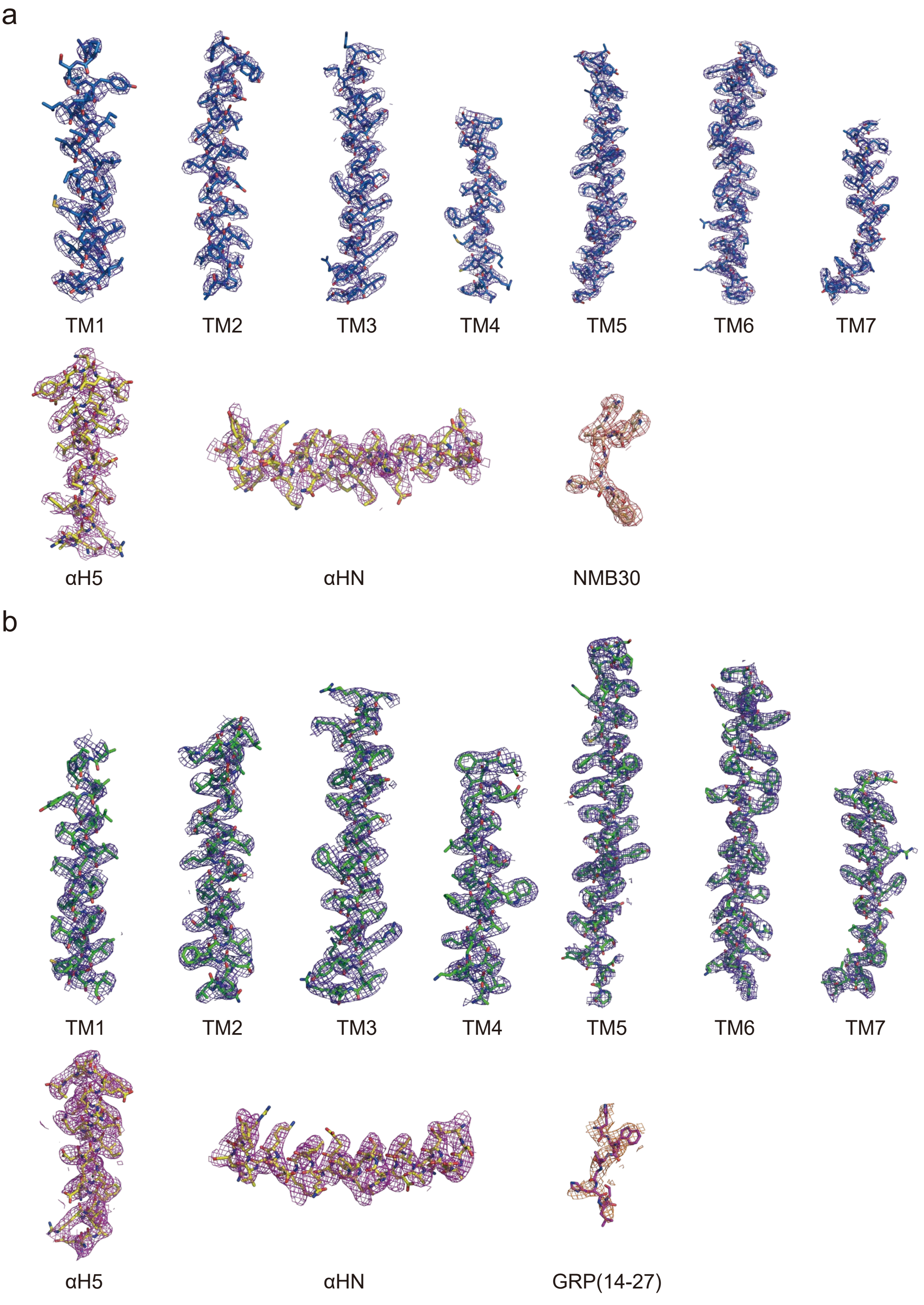
a
TM1
TM2
TM3
TM4
TM5
TM6
TM7
αH5
αHN
NMB30
b
TM1
TM2
TM3
TM4
TM5
TM6
TM7
αH5
αHN
GRP(14-27)

Fig. S6

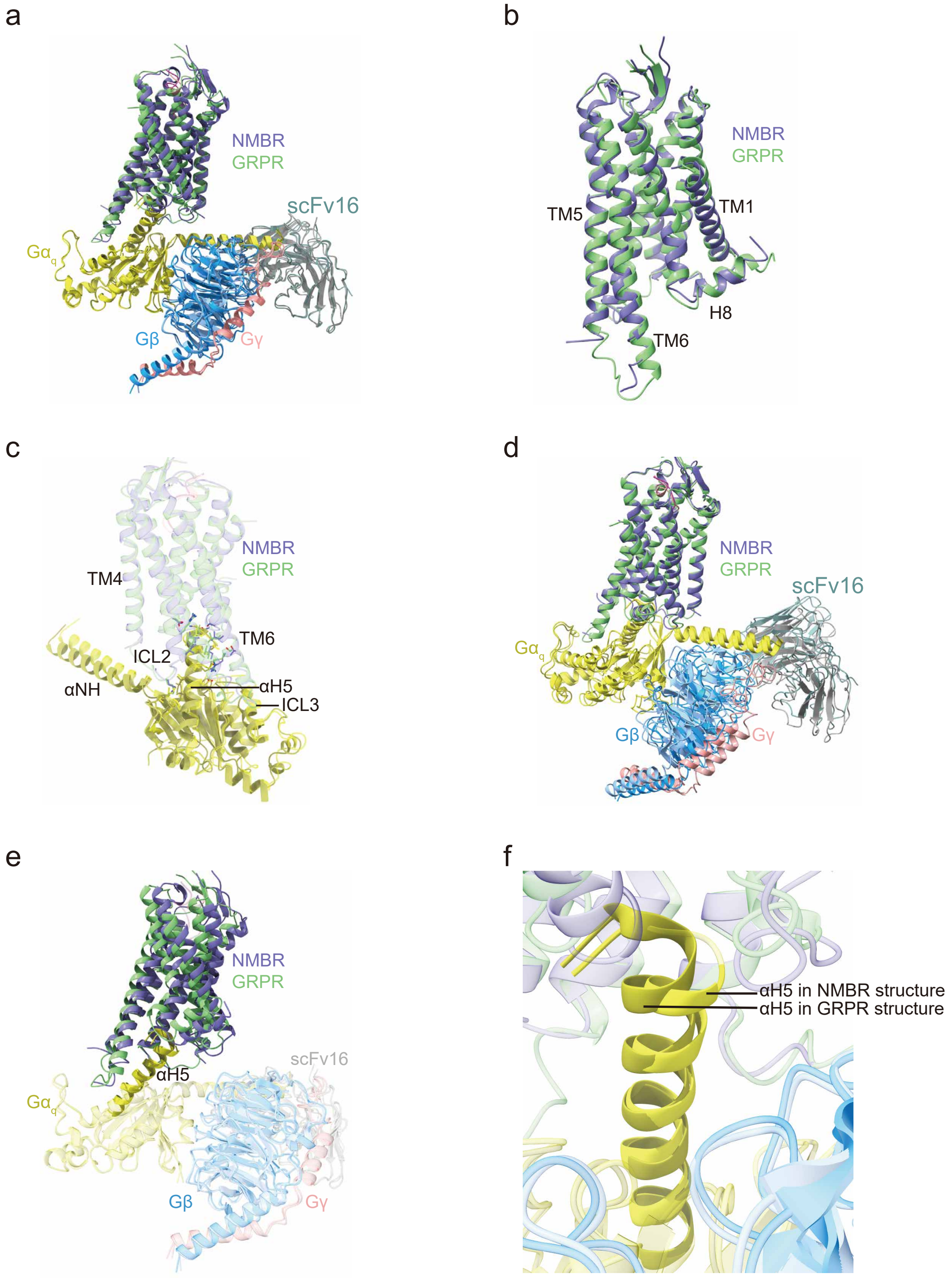
a
NMBR
GRPR
scFv16
Gα$_q$
Gβ
Gγ
b
NMBR
GRPR
TM5
TM1
H8
TM6
c
NMBR
GRPR
TM4
TM6
ICL2
αNH
αH5
ICL3
d
NMBR
GRPR
scFv16
Gα$_q$
Gβ
Gγ
e
NMBR
GRPR
scFv16
αH5
Gα$_q$
Gβ
Gγ
f
αH5 in NMBR structure
αH5 in GRPR structure

Fig. S7

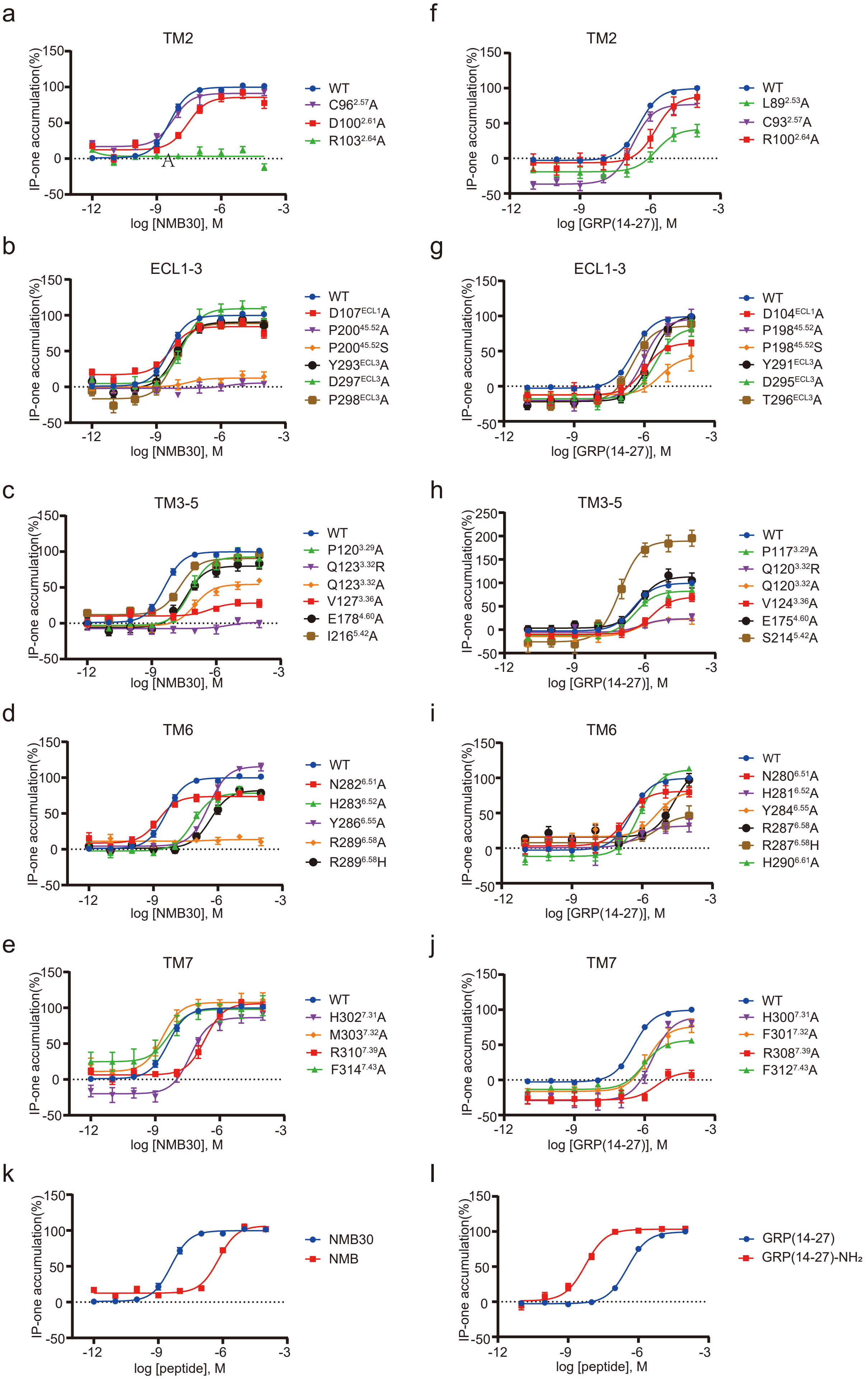

a
TM2
IP-one accumulation(%)
log [NMB30], M
WT
C96$^{2.57}$A
D100$^{2.61}$A
R103$^{2.64}$A
f
TM2
IP-one accumulation(%)
log [GRP(14-27)], M
WT
L89$^{2.53}$A
C93$^{2.57}$A
R100$^{2.64}$A
b
ECL1-3
IP-one accumulation(%)
log [NMB30], M
WT
D107$^{ECL1}$A
P200$^{45.52}$A
P200$^{45.52}$S
Y293$^{ECL3}$A
D297$^{ECL3}$A
P298$^{ECL3}$A
g
ECL1-3
IP-one accumulation(%)
log [GRP(14-27)], M
WT
D104$^{ECL1}$A
P198$^{45.52}$A
P198$^{45.52}$S
Y291$^{ECL3}$A
D295$^{ECL3}$A
T296$^{ECL3}$A
c
TM3-5
IP-one accumulation(%)
log [NMB30], M
WT
P120$^{3.29}$A
Q123$^{3.32}$R
Q123$^{3.32}$A
V127$^{3.36}$A
E178$^{4.60}$A
I216$^{5.42}$A
h
TM3-5
IP-one accumulation(%)
log [GRP(14-27)], M
WT
P117$^{3.29}$A
Q120$^{3.32}$R
Q120$^{3.32}$A
V124$^{3.36}$A
E175$^{4.60}$A
S214$^{5.42}$A
d
TM6
IP-one accumulation(%)
log [NMB30], M
WT
N282$^{6.51}$A
H283$^{6.52}$A
Y286$^{6.55}$A
R289$^{6.58}$A
R289$^{6.58}$H
i
TM6
IP-one accumulation(%)
log [GRP(14-27)], M
WT
N280$^{6.51}$A
H281$^{6.52}$A
Y284$^{6.55}$A
R287$^{6.58}$A
R287$^{6.58}$H
H290$^{6.61}$A
e
TM7
IP-one accumulation(%)
log [NMB30], M
WT
H302$^{7.31}$A
M303$^{7.32}$A
R310$^{7.39}$A
F314$^{7.43}$A
j
TM7
IP-one accumulation(%)
log [GRP(14-27)], M
WT
H300$^{7.31}$A
F301$^{7.32}$A
R308$^{7.39}$A
F312$^{7.43}$A
k
IP-one accumulation(%)
log [peptide], M
NMB30
NMB
l
IP-one accumulation(%)
log [peptide], M
GRP(14-27)
GRP(14-27)-NH$_2$

### Fig. S8

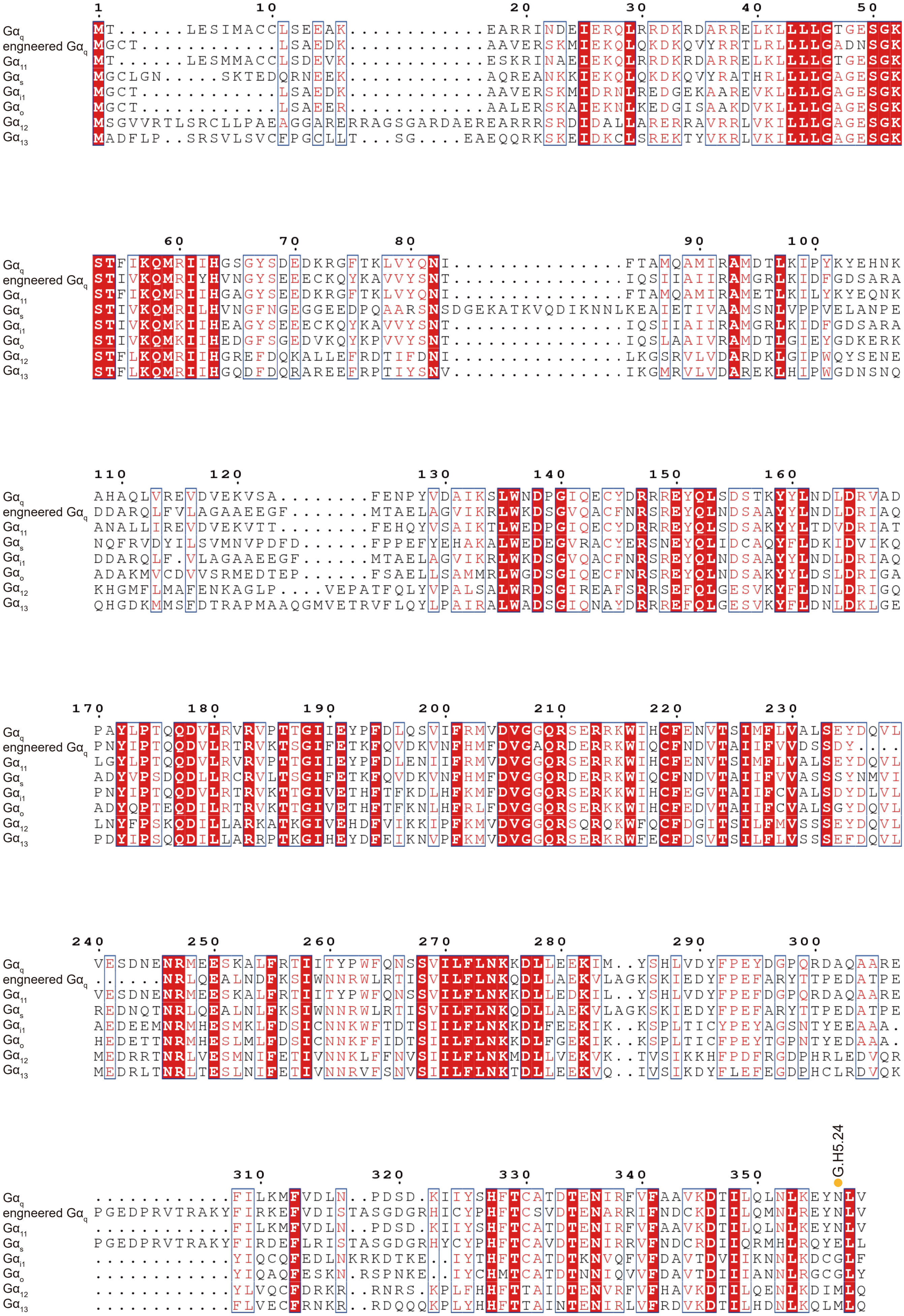
Gαq
engneered Gαq
Gα11
Gαs
Gαi1
Gαo
Gα12
Gα13
1
10
20
30
40
50
60
70
80
90
100
110
120
130
140
150
160
170
180
190
200
210
220
230
240
250
260
270
280
290
300
310
320
330
340
350
G.H5.24

Fig. S9

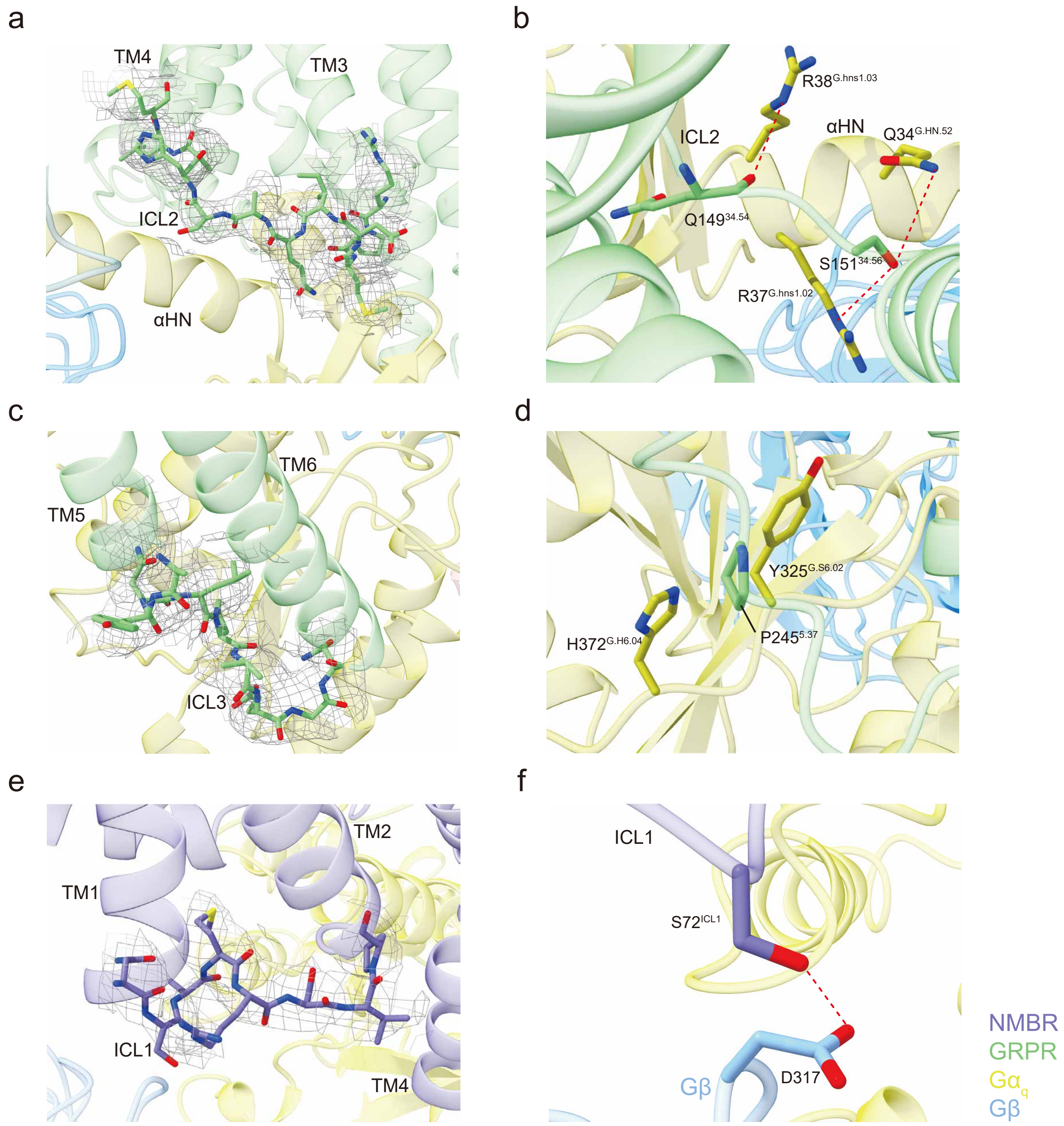
a
TM4
TM3
ICL2
αHN
b
R38$^{G.hns1.03}$
ICL2
αHN
Q34$^{G.HN.52}$
Q149$^{34.54}$
S151$^{34.56}$
R37$^{G.hns1.02}$
c
TM6
TM5
ICL3
d
Y325$^{G.S6.02}$
H372$^{G.H6.04}$
P245$^{5.37}$
e
TM2
TM1
ICL1
TM4
f
ICL1
S72$^{ICL1}$
Gβ
D317
NMBR
GRPR
Gα$_q$
Gβ

Fig. S10

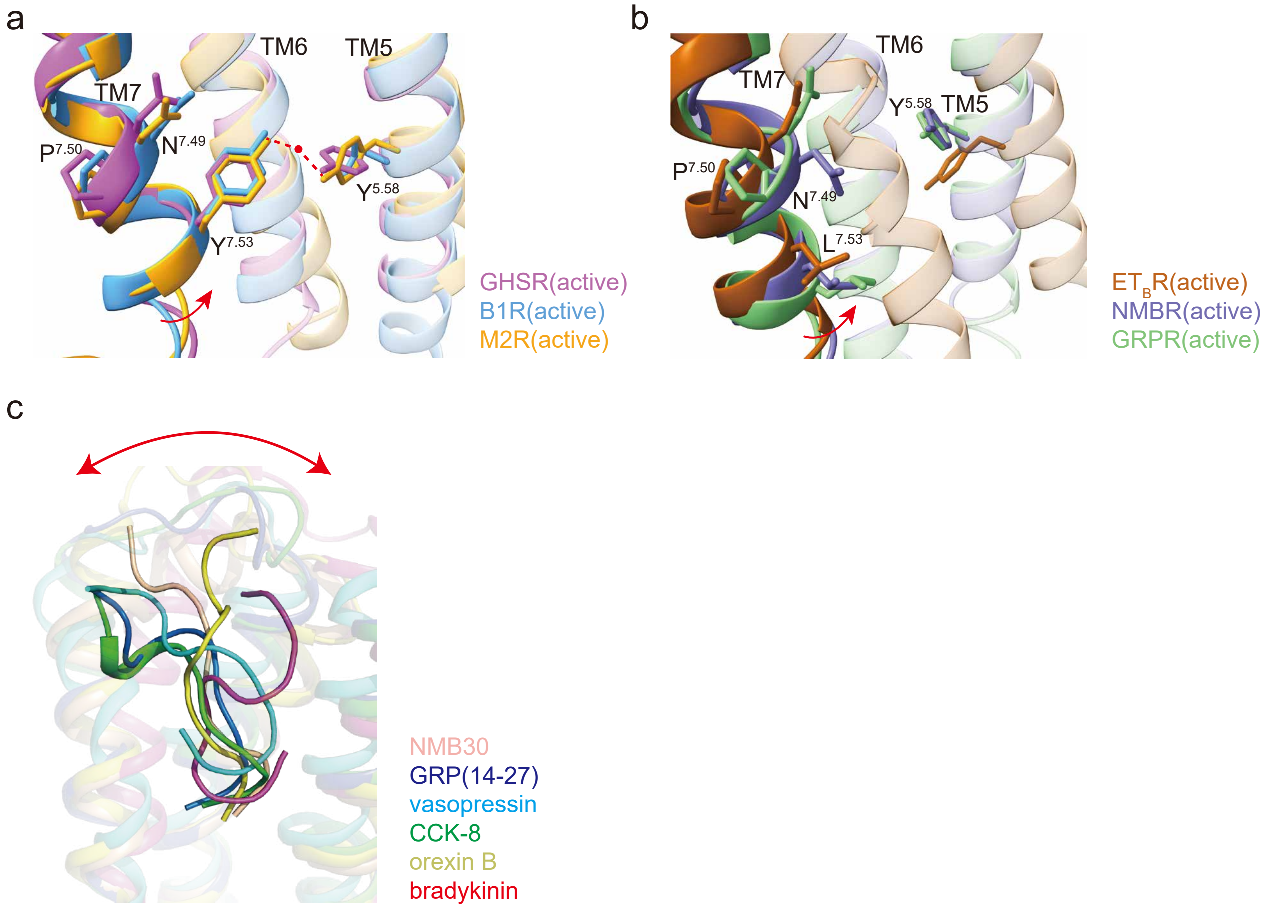
a
TM6
TM5
TM7
P7.50
N7.49
Y7.53
Y5.58
GHSR(active)
B1R(active)
M2R(active)
b
TM6
TM7
Y5.58
TM5
P7.50
N7.49
L7.53
ETBR(active)
NMBR(active)
GRPR(active)
c
NMB30
GRP(14-27)
vasopressin
CCK-8
orexin B
bradykinin

**Table S1 Cryo-EM data collection, model refinement and validation statistics.**

| | NMR30-NMBR-$G_q$ complex | GRP(14-27)-GRPR-$G_q$ complex |
|---|---|---|
| **Data collection and processing** | | |
| Magnification | 81,000 | 96,000 |
| Voltage (kV) | 300 | 300 |
| Electron exposure ($e^-/Å^2$) | 50 | 50 |
| Defocus range (μm) | -1.0~-3.0 | -1.0~-3.0 |
| Pixel size (Å) | 1.04 | 0.8 |
| Symmetry imposed | C1 | C1 |
| Initial particle projections (no.) | 3,034,736 | 3,365,839 |
| Final particle projections (no.) | 355,509 | 245,906 |
| Map resolution (Å) | 3.15 | 3.30 |
| Map resolution range (Å) | 3.0-4.5 | 3.0-4.5 |
| FSC threshold | 0.143 | 0.143 |
| **Model Refinement** | | |
| Refinement package | PHENIX-1.17.1-3660 | PHENIX-1.17.1-3660 |
| Real or reciprocal space | Real space | Real space |
| Model-Map CC (mask) | 0.65 | 0.64 |
| Model resolution (Å) | 3.3 | 3.3 |
| FSC threshold | 0.5 | 0.5 |
| B factors (Å2, mean value) | | |
| Protein residues | 30.37 | 104.85 |
| Ligands | - | 59.28 |
| **Model composition** | | |
| Non-hydrogen atoms | 9,109 | 9,131 |
| Protein residues | 1,163 | 1166 |
| R.m.s. deviations | | |
| Bond lengths (Å) | 0.001 | 0.002 |
| Bond angles (°) | 0.393 | 0.538 |
| **Validation** | | |
| MolProbity score | 1.49 | 1.34 |
| Clashscore | 9.21 | 6.23 |
| Rotamer outliers (%) | 0.20 | 0.41 |
| Ramachandran plot | | |
| Favored (%) | 98.60 | 98.51 |
| Allowed (%) | 1.40 | 1.49 |
| Disallowed (%) | 0 | 0 |
| **Data availability** | | |
| EMDB entry | | |
| PDB entry | | |

**Table S2 Amino acid sequences of Bn-related peptides used or mentioned in this study.**

| Peptide name | Bombesin analogue | N-Terminus | position relative to Bn | | | | | | | | | | | | | |
|---|---|---|---|---|---|---|---|---|---|---|---|---|---|---|---|---|
| | | | 1 | 2 | 3 | 4 | 5 | 6 | 7 | 8 | 9 | 10 | 11 | 12 | 13 | 14 |
| | Bombesin-related | | | | | | | | | | | | | | | |
| Bn | Bombesin | | pE | Q | R | L | G | N | Q | W | A | V | G | H | L | M-NH$_2$ |
| GRP27 | GRP (1–27) | VPLPAGGGTVLTK | M | Y | P | R | G | N | H | W | A | V | G | H | L | M-NH2 |
| GRP (14–27) | GRP (14–27) | | M | Y | P | R | G | N | H | W | A | V | G | H | L | M |
| Aly | Alytesin | | | G | R | L | G | T | Q | W | A | V | G | H | L | M-NH$_2$ |
| NMC | Neuromedin C | | | | | | G | N | H | W | A | V | G | H | L | M-NH2 |
| | Neuromedin B-related | | | | | | | | | | | | | | | |
| NMB | Neuromedin B | | | | | | G | N | L | W | A | T | G | H | F | M-NH$_2$ |
| NMB30 | Neuromedin B (1-30) | LSWDLPEPRSRASKIR | V | H | R | R | G | N | L | W | A | T | G | H | F | M-NH2 |
| Roh-Lit | Rohdei-litorin | | | | | | | pE | L | W | A | T | G | H | F | M-NH$_2$ |
| Lit | Litorin | | | | | | | pE | Q | W | A | V | G | H | F | M-NH2 |
| Ran | Ranatensin | | | | | pE | V | P | Q | W | A | V | G | H | F | M-NH$_2$ |
| | Phyllolitorin-related | | | | | | | | | | | | | | | |
| PLL | Phyllolitorin | | | | | | | pE | L | W | A | V | G | S | F | M-NH2 |
| LeuPLL | [Leu$_8$]phyllolitorin | | | | | | | pE | L | W | A | V | G | S | L | M-NH2 |
| | | | | | | | 1 | 2 | 3 | 4 | 5 | 6 | 7 | 8 | 9 | 10 |
| | | | | | | | position relative to NMB or NMC | | | | | | | | | |

**Table S3 Ligand binding affinities and expression levels of WT and mutated NMBR and GRPR. The wild type (WT) and mutants of NMBR and GRPR discussed in this manuscript were individually analyzed. The affinities are derived from at least 3 independent experiments using IP1 function assay. The expression level of mutant NMBR and GRPR were normalized to wild-type NMBR and GRPR as 100%, respectively. Each data point represents mean ± standard error of the mean (S.E.M.). All data were analyzed by two-sided Student's t test. *P<0.05, **P<0.01, ***P<0.001 vs. WT. Source data are available online. Definitions: NA – not applicable; NT, not tested.**

| Residue Number | NMBR mutant | $pEC_{50}$±S.E.M. | Emax± S.E.M.(%WT) | Expression (% of WT) | GRPR mutant | $pEC_{50}$±S.E.M. | Emax± S.E.M.(%WT) | Expression (% of WT) |
|---|---|---|---|---|---|---|---|---|
| - | WT | 8.44±0.14 | 100 | 100±4.14 | WT | 6.45±0.07 | 100 | 100±29.335 |
| 2.53 | L92A | NT | NT | NT | L89A | 5.97±0.30 | 41.31±18.00 | 155.57±28.22 |
| 2.57 | C96A | 8.25±0.09 | 91.15±8.01 | 74.71±3.79 | C93A | 6.78±0.07* | 135.42±62.17 | 262.60±20.67 |
| 2.61 | D100A | 7.58±0.13** | 85.81±12.88 | 53.52±7.35 | D97A | NT | NT | NT |
| 2.64 | R103A | NA | NA | 90.69±3.02 | R100A | 5.72±0.11** | 88.66±25.73 | 170.49±22.41 |
| ECL1 | D107A | 8.44±0.16 | 84.17±10.41 | 68.84±3.05 | D104A | 5.87±0.26 | 63.70±1.46** | 85.32±7.97 |
| 3.29 | P120A | 7.21±0.04*** | 92.66±20.11 | 120.21±7.57 | P117A | 6.37±0.21 | 128.81±59.52 | 346.31±20.86 |
| 3.32 | Q123R | NA | NA | 40.12±3.99 | Q120R | NA | NA | NT |
| 3.32 | Q123A | 7.05±0.14*** | 54.21±5.36* | 7.15±0.24 | Q120A | NA | NA | 13.09±3.43 |
| 3.36 | V127A | NA | NA | 24.72±5.30 | V124A | 5.65±0.16* | 94.37±34.90 | 184.24±25.85 |
| 3.43 | L134Q | 7.70±0.15* | 38.33±9.00* | 11.81±0.73 | L131Q | NA | NA | 128.07±58.55 |
| 4.60 | E178A | 7.43±0.18** | 80.05±13.17 | 73.16±1.90 | E175A | 6.28±0.08 | 113.07±16.97 | 155.24±5.12 |
| 45.52 | P200A | NA | NA | 59.29±1.05 | P198A | 6.00±0.09* | 96.52±26.70 | 156.74±16.41 |
| 45.52 | P200S | NA | NA | 43.96±7.55 | P198S | 5.31±0.07*** | 45.27±39.47 | 94.46±6.80 |
| 5.42 | I216A | 7.70±0.03** | 90.37±6.84 | 14.71±1.51 | S214A | 7.08±0.09** | 189.80±29.77 | 185.15±21.86 |
| 6.51 | N282A | 8.76±0.08 | 73.85±4.13* | 83.74±10.51 | N280A | 6.53±0.36 | 82.13±9.85 | 149.76±6.71 |
| 6.52 | H283A | 7.14±0.14*** | 78.53±4.91* | 48.61±6.28 | H281A | 6.02±0.22 | 32.65±9.35* | 211.66±12.67 |
| 6.55 | Y286A | 6.42±0.07*** | 116.03±7.01 | 47.01±3.35 | Y284A | 5.46±0.25 | 79.50±1.73** | NT |
| 6.58 | R289A | NA | NA | 4.81±4.97 | R287A | NA | NA | 130.96±24.39 |
| 6.58 | R289H | 6.30±0.13*** | 82.99±2.61* | 98.05±6.46 | R287H | 5.27±0.09*** | 47.19±25.23 | 43.88±17.99 |
| 6.61 | N292 | NT | NT | 81.52±6.00 | H290A | 6.05±0.15 | 144.20±27.65 | 269.90±18.37 |
| ECL3 | Y293A | 8.15±0.12 | 90.39±15.12 | 122.45±4.60 | Y291A | 5.72±0.21 | 101.72±3.61 | 260.51±74.72 |
| ECL3 | D297A | 7.80±0.20 | 109.32±16.48 | 24.16±0.70 | D295A | 5.49±0.13** | 83.46±21.40 | 123.10±17.66 |
| ECL3 | P298A | 8.21±0.15 | 88.68±10.42 | 42.75±6.08 | T296A | 6.40±0.27 | 121.95±44.58 | 275.41±15.80 |
| 7.31 | H302A | 7.37±0.13* | 86.72±13.19 | 48.07±1.78 | H300A | 5.62±0.08*** | 124.90±23.70 | 145.88±30.42 |
| 7.32 | M303A | 8.65±0.03 | 107.57±13.70 | 19.15±1.44 | F301A | 5.71±0.25 | 150.32±50.02 | 77.74±7.57 |
| 7.39 | R310A | 6.69±0.15*** | 106.12±8.98 | 37.09±2.85 | R308A | 5.52±0.12** | 10.90±10.26* | 214.81±35.21 |
| 7.43 | F314A | 8.37±0.05 | 98.02±18.40 | 64.70±4.36 | F312A | 6.03±0.03*** | 74.81±15.01 | 85.97±10.67 |
| - | NMB | 6.21±0.06*** | 106.55±5.03 | 100±4.14 | GRP(14-27)-NH2 | 8.30±0.12*** | 103.10±1.54 | 100±29.335 |

**Table S4 Sequence alignment of the key residues in sodium site, DRY motif, PV(I)F motif, toggle switch and NPxxY motif, as well as residues involved in disulfide bond formation in bombesin receptors.**

| | Sodium site | | Disulfide bond | | | DRY motif | | | PV(I)F motif | | | toggle switch | | | NPxxY motif | | |
|---|---|---|---|---|---|---|---|---|---|---|---|---|---|---|---|---|---|
| Residue position | 2.50 | 3.39 | 3.25 | 4.64 | 5.34 | 3.49 | 3.50 | 3.51 | 5.50 | 3.40 | 6.44 | 6.44 | 6.48 | 7.43 | 7.49 | 7.50 | 7.53 |
| NMBR | D | S | C | T | H | D | R | Y | P | V | F | F | W | F | N | P | L |
| GRPR | D | S | C | S | H | D | R | Y | P | V | F | F | W | F | N | P | L |
| BRS3 | D | S | C | S | L | D | R | Y | P | V | F | F | W | F | N | P | L |
| Class A conserved | D | S | C | x | x | D | R | Y | P | I | F | F | W | F/Y | N | P | Y |
